# The calcineurin pathway regulates extreme thermotolerance, cell membrane and wall integrity, antifungal resistance, and virulence in *Candida auris*

**DOI:** 10.1101/2025.01.02.631159

**Authors:** Hyunjin Cha, Doyeon Won, Seun Kang, Eui-Seong Kim, Kyung-Ah Lee, Won-Jae Lee, Kyung-Tae Lee, Yong-Sun Bahn

**Affiliations:** Department of Biotechnology, College of Life Science and Biotechnology, Yonsei University, Seoul, Republic of Korea; Korea Zoonosis Research Institute, Jeonbuk National University, Iksan, Jeonbuk, Republic of Korea; School of Biological Sciences, Seoul National University, Seoul 08826, South Korea; Saeloun Bio Inc., Seoul 08826, South Korea

**Keywords:** *C. auris*, Calcineurin, Cna1, Cnb1, Crz1, Multidrug resistance

## Abstract

*Candida auris*, an emerging fungal pathogen characterized by its multidrug resistance and high mortality rates, poses a significant public health challenge. Despite its importance, the signaling pathways governing virulence and antifungal resistance in *C. auris* remain poorly understood. This study investigates the calcineurin pathway in *C. auris*, critical for virulence and antifungal resistance in other fungal pathogens. Calcineurin, a calcium/calmodulin-dependent protein phosphatase, comprises a catalytic subunit (Cna1) and a regulatory subunit (Cnb1) in *C. auris*. Our findings reveal that deletion of *CNA1* or *CNB1* disrupts extreme thermotolerance and cell membrane and wall integrity, leading to increased susceptibility to azoles and echinocandins. Moreover, we identified a downstream transcription factor, Crz1, which plays a central role in this pathway in other fungal species. Deletion of *CRZ1* resulted in similar membrane integrity defects observed in the *cna1*Δ and *cnb1*Δ mutants and increased susceptibility to azole drugs. Supporting it, fluconazole treatment induced Crz1 nuclear translocation in a Cna1-dependent manner. However, unlike *cna1*Δ and *cnb1*Δ mutants, the *crz1*Δ mutant displayed increased resistance to echinocandins, suggesting the opposing roles for Crz1 in regulating cell wall integrity. Nevertheless, echinocandins also promoted Crz1 nuclear translocation via Cna1, underscoring the complex regulatory mechanisms at play. Cna1 was found to be required for virulence in both the *Drosophila* systemic infection model and the murine skin infection model. However, in a systemic murine infection model, both calcineurin and Crz1 appeared dispensable for *C. auris* virulence. Our findings highlight that the evolutionarily conserved calcineurin pathway employs distinct regulatory mechanisms to perform divergent roles in regulating cell wall and membrane integrity, antifungal drug resistance, and virulence in *C. auris*.

**Author Summary:** The fungal pathogen *Candida auris* presents a global health threat due to its multidrug resistance and high mortality rates. Despite its clinical significance, the molecular mechanisms underlying its virulence and antifungal resistance remain poorly understood. This study investigates the complex role of the calcineurin signaling pathway in *C. auris* pathogenicity. Deletion of the calcineurin complex impairs extreme thermotolerance and compromises cell membrane and wall integrity, leading to increased susceptibility to azoles and echinocandins, antifungal agents targeting the cell membrane and wall, respectively. We also identified the Crz1 transcription factor as a downstream target of calcineurin signaling. Interestingly, unlike calcineurin mutants, Crz1 mutants are susceptible only to cell-membrane-targeting azoles but surprisingly exhibit increased resistance to cell-wall-targeting echinocandins, suggesting that the calcineurin pathway may regulate multiple transcription factors, including Crz1. Additionally, calcineurin was found to be essential for virulence in vivo, using two different animal infection models, Drosophila and mice. These findings highlight the essential role of the calcineurin pathway in *C. auris* virulence, offering novel insights into its role in antifungal resistance and virulence, and paving the way for targeted therapies.

## Introduction

Fungal infections, particularly those caused by *Candida* species, pose a significant global health threat, accounting for over 3.75 million deaths annually (1). The fungal pathogen *Candida auris* has emerged as a particular concern because it often affects immunocompromised patients and is associated with high mortality rates (2). *C. auris* often shows resistance to several antifungal drugs, including azoles, amphotericin B, and echinocandins, which are commonly used for candidiasis treatment. In 2020, the CDC reported that 86% of *C. auris* isolates in the United States are resistant to azoles, 26% to amphotericin B, and about 1% to echinocandins (3). Due to these factors, the World Health Organization (WHO) has classified *C. auris* as a critical priority fungal pathogen. Consequently, understanding its mechanisms of virulence and antifungal resistance is crucial for developing new antifungal therapies.

The calcineurin pathway is an evolutionarily conserved eukaryotic signaling cascade. Calcineurin is a serine/threonine protein phosphatase composed of catalytic (Cna1) and regulatory (Cnb1) subunits. It is activated when the calcium ions (Ca^2+^) enter the cytosol through membrane transporters. The influx of Ca^2+^ is recognized by calmodulin, leading to the formation of the Ca^2+^-calmodulin complex. The cytosolic Ca^2+^ also binds to Cnb1, altering its conformation of Cna1 and allowing calcineurin to interact with the Ca^2+^/calmodulin complex for its activation (4). Following activation, calcineurin dephosphorylates a downstream transcriptional factor. This factor then translocates from the cytosol to the nucleus, regulating the expression of calcineurin-dependent genes (4). In humans, calcineurin targets NFAT (nuclear factor of activated T cells), while in yeast, the target is Crz1 (calcineurin-responsive zinc finger 1) (5).

The calcineurin-Crz1 pathway plays a critical role in environmental adaption for yeast and various fungal pathogens. In *Saccharomyces cerevisiae*, calcineurin is indispensable for responding to high concentrations of cations (6). Furthermore, the deletion of *CRZ1* results in phenotypes similar to those of calcineurin mutants, though less severe (7). In *Candida albicans*, the calcineurin pathway is crucial for regulating thermotolerance, cell wall integrity, serum survival, virulence, and drug tolerance (4). Similarly, in *Candida glabrata*, it influences thermotolerance, intracellular architecture, and virulence (8). In *Cryptococcus neoformans*, calcineurin is essential for stress adaptation and virulence, particularly under high temperature conditions, impacting cell wall remodeling, calcium transport, and pheromone production (6). Additionally, in *C. neoformans*, substrates other than Crz1 have been identified as targets of calcineurin (9).

Despite the conserved and divergent roles of the calcineurin pathway in the growth, development, antifungal drug resistance, and pathogenicity across other fungal pathogens, its pathobiological functions and regulatory mechanism remain unexplored in *C. auris*. In this study, we identified and functionally characterized the catalytic and regulatory subunits of calcineurin in *C. auris*. Here, we demonstrate that the calcineurin pathway is required for maintaining cell membrane and wall integrity, affecting resistance to azoles and echinocandins, and virulence in *C. auris*. We also discovered that Crz1 is one of the calcineurin targets but plays overlapping and distinct roles with calcineurin. Collectively, our study highlights that the calcineurin pathway plays conserved and distinct pathobiological roles with distinct regulatory mechanisms in this emerging multidrug-resistant fungal pathogen.

## Results

### Calcineurin is essential for extreme thermotolerance and the maintenance of cell membrane and wall integrity in *C. auris*

To identify the catalytic (Cna1) and regulatory (Cnb1) subunits of calcineurin in *C. auris*, we queried the *Candida* genome database. Orthologs were identified in the clade I *C. auris* wild-type strain (AR0387) as Cna1 (B9J08_004950) and Cnb1 (B9J08_001732). Sequence analysis revealed that both subunits are highly conserved across eukaryotic species (S1A and S1B Fig). Protein domain analysis using the InterPro database (https://www.ebi.ac.uk/interpro/) showed that Cna1 harbors a conserved serine/threonine-specific protein phosphatase domain, which catalyzes the dephosphorylation of phosphoserine and phosphothreonine residues. The size of Cna1 is comparable to its orthologs in other fungal species (Fig 1A). Cnb1 was found to contain two EF-hand motifs, characteristic calcium-binding domains (Fig 1A). Structural modeling using AlphaFold3 predicted a heterodimeric calcineurin complex in *C. auris* consisting of Cna1 and Cnb1, featuring helix-loop-helix structures (Fig 1B). To investigate the roles of calcineurin in *C. auris*, we generated *cna1*Δ, *cnb1*Δ, and *cnb1*Δ *cna1*Δ mutants in the clade I AR0387 (B8441) strain background (S2A, S2B, and S2C Fig). To confirm the phenotypic features of the mutants, we also constructed complemented strains by integrating the wild-type allele into its original locus (*cna1*Δ::*CNA1*, and *cnb1*Δ::*CNB1*) (S2D and S2E Fig). Under nutrient-rich growth conditions at ambient temperatures (25°C to 30°C), the calcineurin mutants exhibited no growth defects (Fig 1C), indicating that the calcineurin pathway is not essential for the normal growth of *C. auris*. As the calcineurin pathway has been implicated in thermotolerance, we assessed the growth of the mutants at elevated temperatures (37°C to 42°C). The *cna1*Δ, *cnb1*Δ, and *cnb1*Δ *cna1*Δ mutants showed no growth defects at high temperatures within the physiological range (37-39°C) (Fig 1C and D), suggesting that the calcineurin pathway is largely dispensable for thermotolerance of *C. auris* under these conditions. However, under severe heat stress conditions exceeding 42°C (43-45°C), *cna1*Δ, *cnb1*Δ, and *cnb1*Δ *cna1*Δ mutants exhibited significant growth defects (Fig 1E and 1F), indicating the critical role of calcineurin in regulating growth at extreme temperatures.

**Fig 1.**
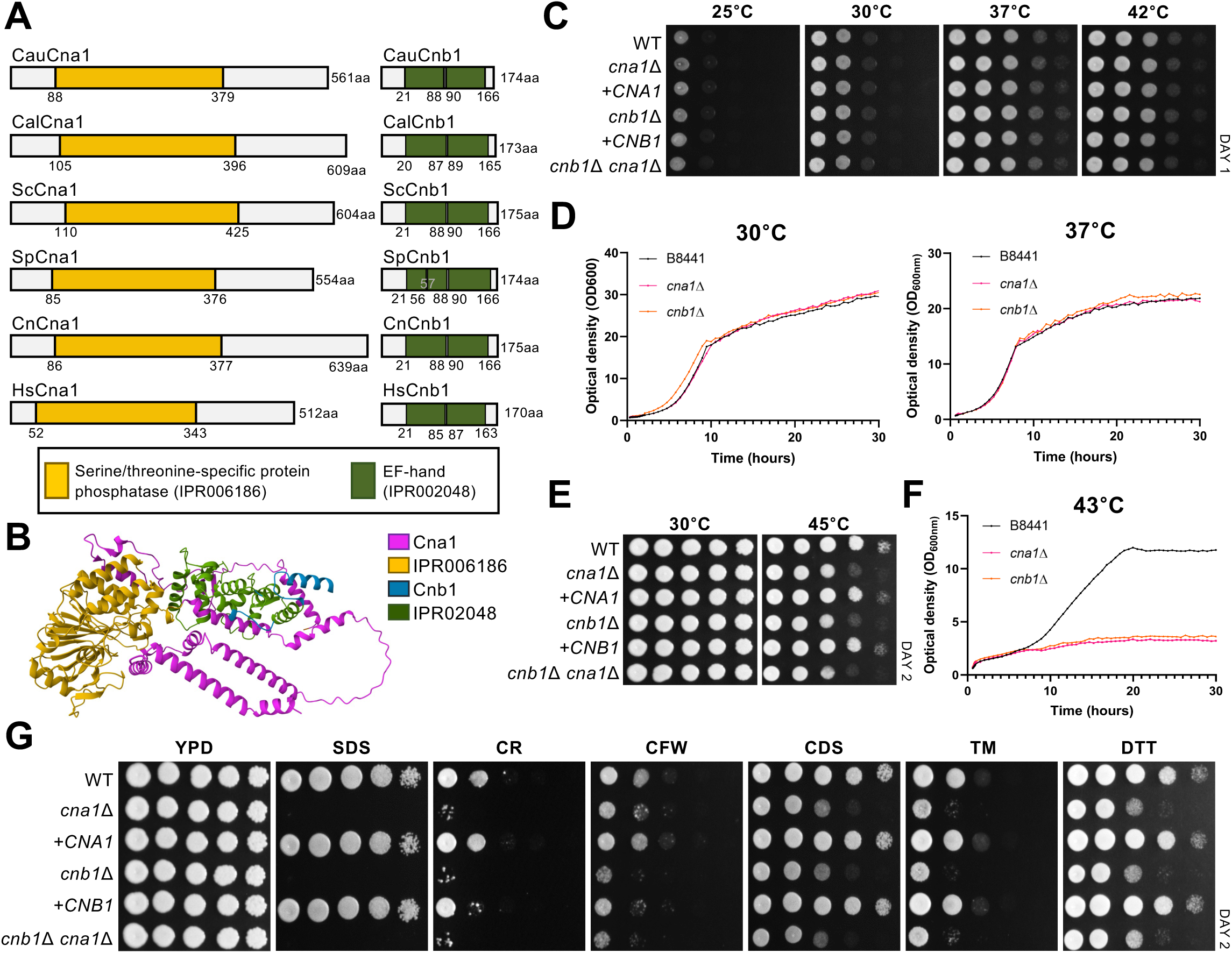
Identification of the calcineurin complex and its roles in growth, thermotolerance, and stress responses in *C. auris*. (A) Domain analysis of Cna1 and Cnb1 in *Candida auris* (Cau), *Candida albicans* (Cal), *Saccharomyces cerevisiae* (Sc), *Schizosaccharomyces pombe* (Sp), *Cryptococcus neoformans* (Cn), and *Homo sapiens* (Hs) was performed using InterPro (https://www.ebi.ac.uk/interpro/). (B) Predicted structures of *C. auris* Cna1 and Cnb1 were modeled using AlphaFold 3. The protein sequences were processed in the order of Cna1 and Cnb1. The color coding is as follows: Cna1 in pink, the IPR006186 domain of Cna1 in yellow, Cnb1 in blue, and the IPR02048 domain of Cnb1 in green. (C) Qualitative spot assays display the growth of the following strains under varying temperatures: wild-type (WT; B8441), *cna1*Δ (YSBA99), *cna1*Δ::*CNA1* (+*CNA1*; YSBA110), *cnb1*Δ (YSBA102), *cnb1*Δ::*CNB1* (+*CNB1*; YSBA113), and *cnb1*Δ *cna1*Δ (YSBA172). WT and mutant strains were cultured overnight in liquid YPD medium at 30°C, serially diluted 10-fold, and spotted on YPD agar plates. Plates were incubated for 1 day at 25°C, 30°C, 37°C, or 42°C. (D) Quantitative growth rates of the WT, *cna1*Δ, and *cnb1*Δ strains were measured at 30°C and 37°C using a multichannel bioreactor (Biosan Laboratories, Inc., Warren, MI) by measuring the OD_600_ for 30 h. (E) WT and mutant strains were spotted on YPD plates and incubated at 30°C or 45°C for 2 days. (F) Growth rates of the WT and mutant strains were monitored at 43°C. (G) WT and mutant strains were spotted on YPD medium supplemented with stress-inducing agents, including 0.1% sodium dodecyl sulfate (SDS), 0.05% Congo red (CR), 0.12 mg/ml calcofluor white (CFW), 250 mM CdSO_4_, 2.5 μg/ml tunicamycin (TM), or 22 mM dithiothreitol (DTT). Plates were incubated for 2 days at 30°C.

We next investigated whether the calcineurin pathway is required for the growth of *C. auris* under various environmental stress conditions, including osmotic stress, oxidative stress, genotoxic stress, and cell membrane/wall-damaging stress (Fig 1G and S3 Fig). Among these stressors, the *cna1*Δ, *cnb1*Δ, and *cnb1*Δ *cna1*Δ mutants demonstrated markedly enhanced sensitivity to cell membrane-destabilizing agents [sodium dodecyl sulfate (SDS)] and cell wall damaging agents [dithiothreitol (DTT), tunicamycin (TM), Congo red (CR), calcofluor white (CFW), and CdSO_4_ (CDS)] (Fig 1G). SDS induces cell membrane stress by disrupting lipid bilayers (10), while CR binds specifically to polysaccharide components such as chitin and glucans in the fungal cell wall (11). DTT and TM, as ER stressors, cause protein misfolding, leading to compromised cell wall integrity (12). CdSO_4_, a toxic heavy metal, interferes with the synthesis and structure of fungal cell wall components such as chitin and glucans, thereby disrupting cell wall integrity (13). Notably, deletion of both *CNA1* and *CNB1* resulted in phenotypes identical to those observed with individual deletions of *CNA1* or *CNB1*, suggesting that Cna1 and Cnb1 function as components of a single calcineurin complex, as expected. In summary, our findings demonstrate that the calcineurin pathway plays a crucial role in maintaining the cell membrane and wall integrity in *C. auris*.

### Calcineurin promotes resistance to azoles and echinocandins in *C. auris*

Given the role of calcineurin in maintaining cell membrane and wall integrity, we investigated its contribution to resistance against clinically relevant antifungal drugs, which target either the fungal cell membrane (azoles and polyenes) or cell wall (echinocandins). We evaluated the susceptibility of the mutants (*cna1*Δ, *cnb1*Δ, and *cnb1*Δ *cna1*Δ) and their complemented strains to echinocandins [caspofungin (CAF), micafungin (MIF), and anidulafungin (ANF)], polyenes [amphotericin B (AMB)], and azoles [fluconazole (FLC), posaconazole (PSC), voriconazole (VRC), ketoconazole (KTC), and itraconazole (ITC)] (Fig 2A). The *cna1*Δ, *cnb1*Δ, and *cnb1*Δ *cna1*Δ mutants exhibited substantially increased susceptibility to all tested echinocandins and azoles compared to the wild-type strain (Fig 2A). The minimum inhibitory concentrations (MICs) of CAF, MIF, and ANF for the wild-type strain were > 80 μg/ml, 1.25 μg/ml, and 1.25 ug/ml, respectively (Fig 2B and S4A Fig). In contrast, the MICs for *cna1*Δ and *cnb1*Δ mutants were markedly reduced: 5 μg/ml for CAF, 0.625 μg/ml for MIF, and 0.625 μg/ml for ANF (Fig 2B and S4A Fig). Consistent with previous reports (14), azoles exhibited fungistatic activity against wild-type *C. auris* and did not completely inhibit its growth (Fig 2B). However, in the absence of calcineurin (*cna1*Δ, *cnb1*Δ, and *cnb1*Δ *cna1*Δ), azoles displayed fungicidal activity (Fig 2B).

**Fig 2.**
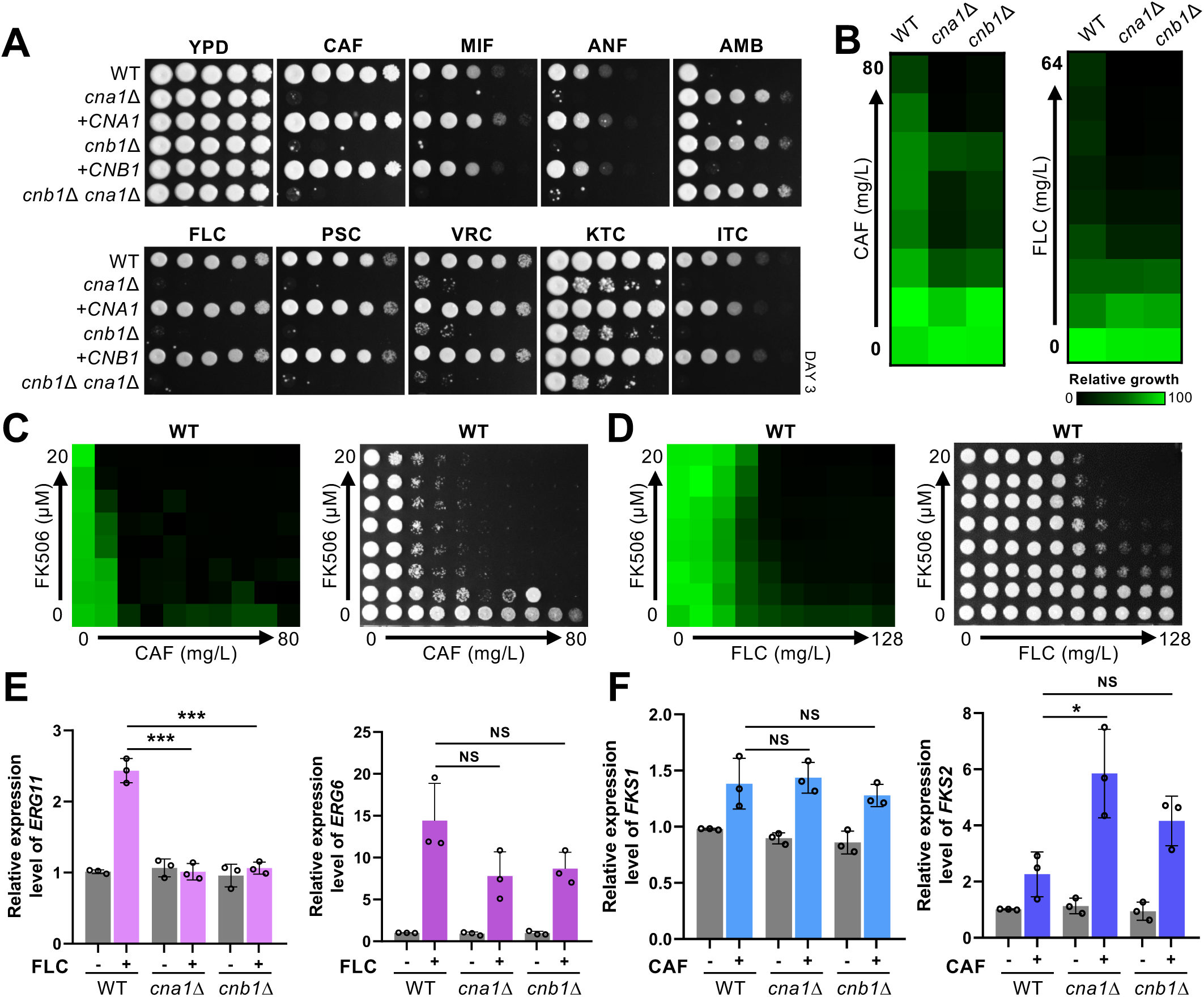
Functions of calcineurin in antifungal resistance in *C. auris*. (A) Qualitative spot assays showing the stress susceptibility of the following strains: WT (B8441), *cna1*Δ (YSBA99), +*CNA1* (YSBA110), *cnb1*Δ (YSBA102), +*CNB1* (YSBA111), and *cnb1*Δ *cna1*Δ (YSBA172). WT and mutant strains were cultured overnight in liquid YPD medium at 30°C, serially diluted 10-fold, and spotted on YPD agar medium supplemented with stressors, including 2.5 μg/ml caspofungin (CAF), 0.15 μg/ml micafungin (MIF), 1 μg/ml anidulafungin (ANF), 3 μg/ml amphotericin B (AMB), 150 μg/ml fluconazole (FLC), 0.5 μg/ml posaconazole (PSC), 0.8 μg/ml voriconazole (VRC), 5 μg/ml ketoconazole (KTC), or 1 μg/ml itraconazole (ITC). Plates were incubated at 30°C for 3 days. (B) EUCAST MIC test results of CAF and FLC in the WT, *cna1*Δ, and *cnb1*Δ strains. The WT strain was grown in YPD medium overnight, washed twice with H_2_O, resuspended in H_2_O, and added (100 μl) to 10 ml of liquid RPMI 1640 medium, which was subsequently loaded into 96-well plates. Drugs serially diluted 2-fold from the indicated concentrations were added to each well. (C) Checkerboard assay results of CAF with FK506 in the WT strain. After growth measurement, cultures were spotted onto YPD agar plates and incubated at 30°C for 24 h to evaluate fungicidal activity. (D) Checkerboard assay results of FLC with FK506 in the WT strain. (E) qRT-PCR analysis of *ERG11* and *ERG6* in the WT, *cna1*Δ, and *cnb1*Δ strains. The strains were cultured for 24 h in the presence of FLC (200 μg/ml) at 30°C in a shaking incubator. Gene expression levels were normalized to *ACT1*, and fold changes were calculated relative to the basal expression level of *ERG11* in WT and mutant strains. Statistical significance was determined using one-way ANOVA with Bonferroni’s multiple-comparison test (*, *P* < 0.05; **, *P* < 0.01; ***, *P* < 0.001; ****, *P* < 0.0001; NS, not significant). (F) qRT-PCR analysis of *FKS1* and *FKS2* in WT, *cna1*Δ, and *cnb1*Δ strains. The strains were cultured overnight at 30°C shaking incubator, sub-cultured to OD_600_ 0.8 in fresh YPD liquid medium, and then cultured for 24 hours with CAF (5 μg/ml) at 30°C shaking incubator.

The observation that the calcineurin pathway promotes resistance to azoles and echinocandins in *C. auris* suggests that calcineurin inhibitors, such as FK506 (tacrolimus) and cyclosporin A, may exhibit synergistic antifungal activity when combined with these drugs. To test this hypothesis, we performed checkerboard assays with FK506, cyclosporin A, azoles, and echinocandins. As anticipated, given our findings that deletion of *CNA1* and *CNB1* does not impair the growth of *C. auris* under basal conditions, FK506 and cyclosporin A alone displayed no antifungal activity at concentrations up to 20 μg/ml (Fig 2C and S4B Fig). However, when combined with echinocandins, both FK506 and cyclosporin A demonstrated fungicidal antifungal activity. Assuming an MIC of 20 μg/ml for FK506 and cyclosporin A, the fractional inhibitory concentration (FIC) index for these combinations with echinocandins were consistently less than 0.5 (Fig 2C and S4B-C Fig), indicating a synergistic relationship. Similarly, FK506 and cyclosporin A exhibited synergistic fungicidal antifungal activity when combined with azoles (Fig 2D and S5 Fig).

Unexpectedly, we found that the *cna1*Δ, *cnb1*Δ, and *cnb1*Δ *cna1*Δ mutants exhibited high resistance to amphotericin B, in stark contrast to their heightened susceptibility to azoles (Fig 2A). Azoles inhibit ergosterol synthesis by targeting lanosterol 14α-demethylase (Erg11), while amphotericin B binds to and sequesters membrane ergosterol. A possible explanation for this finding is that the calcineurin pathway may regulate *ERG11* expression in *C. auris*, as reduced membrane ergosterol could confer resistance to amphotericin B. To explore this hypothesis, we assessed basal *ERG11* expression levels, which were comparable between the wild-type and calcineurin mutant strains (Fig 2E). In response to FLC treatment, *ERG11* expression was upregulated in the wild type, possibly as a compensatory mechanism. However, this induction was absent in the *cna1*Δ and *cnb1*Δ mutants (Fig 2E), suggesting that the calcineurin pathway plays a role in the transcriptional regulation of ergosterol biosynthetic genes.

We further investigated echinocandin resistance, which is associated with altered expression of *FKS1* and *FKS2*, encoding β-1,3-glucan synthase, the target of echinocandins. Basal and echinocandin-induced expression levels of *FKS1* in the wild type, *cna1*Δ, and *cnb1*Δ mutants, revealing no significant changes in *FKS1* expression (Fig 2F). In contrast, *FKS2* expression was markedly upregulated in the *cna1*Δ mutant (Fig 2F), suggesting that echinocandin resistance in *C. auris* may be linked to the upregulation of *FKS2* rather than *FKS1*, particularly in the absence of Cna1. Collectively, our findings demonstrated that the calcineurin pathway is critical for promoting resistance to azoles and echinocandins in *C. auris*.

### Conserved roles of calcineurin in other *C. auris* clades

*Candida auris* is classified into five clades according to the isolated location: clade I (South Asian), clade II (East Asian), clade III (African), clade IV (South American), and clade V (Iran) (15). Each clade exhibits distinct genotypic and phenotypic variations (16, 17). To determine whether the role of the calcineurin pathway is conserved among different clades of *C. auris*, we performed a phylogenetic analysis, which revealed that Cna1 is highly conserved across all clades (Fig 3A).

**Fig 3.**
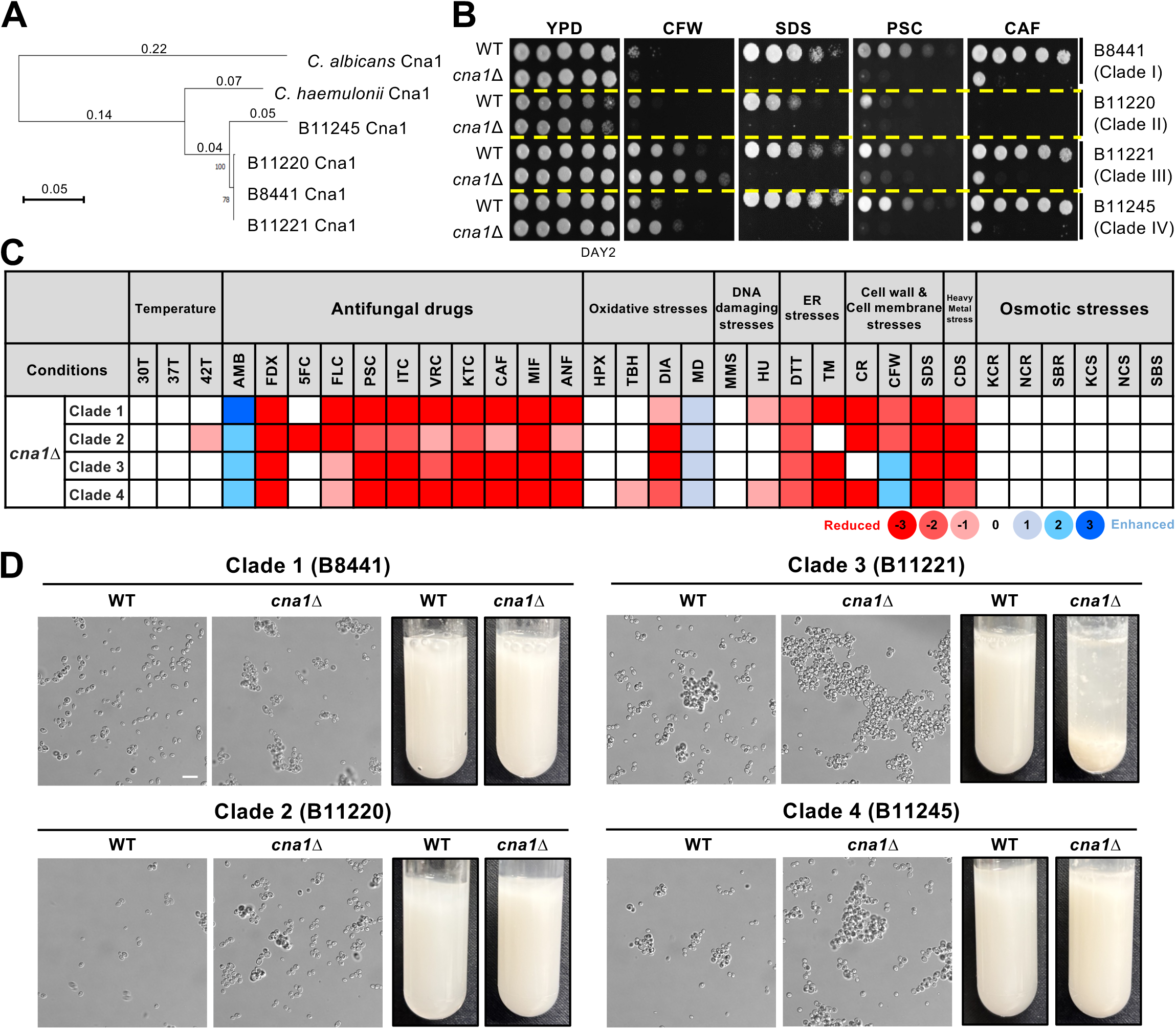
Functional roles of calcineurin across *C. auris* clades. (A) Phylogenetic analysis of Cna1 orthologs across *C. albicans*, *C. haemulonii*, and four *C. auris* clades. (B) Qualitative spot assays showing the stress susceptibility of clade I WT (B8441), clade I *cna1*Δ (YSBA99), clade II WT (B11220), clade II *cna1*Δ (YSBA332), clade III WT (B11221), clade III *cna1*Δ (YSBA336), clade IV WT (B11245), and clade IV *cna1*Δ (YSBA365) strains. WT and mutant strains were cultured overnight in liquid YPD medium at 30°C, serially diluted 10-fold, and spotted on YPD medium supplemented with stressors, including 0.12 mg/ml calcofluor-white, 0.2% sodium dodecyl sulfate (SDS), 0.05 μg/ml PSC, or 0.1 μg/ml CAF. (C) Phenome heat map of *cna1*Δ mutants across four different *C. auris* clades. Phenotype scores are color-coded based on qualitative or semi-quantitative measurements under the indicated growth conditions. Abbreviations: 30T, 30°C; 37T, 37°C; 42T, 42°C; AMB, amphotericin B; FDX, fludioxonil; 5FC, 5-flucytosine; FLC, fluconazole; PSC, posaconazole; ITC, itraconazole; VRC, voriconazole; KTC, ketoconazole; CAF, caspofungin; MIF, micafungin; ANF, anidulafungin; HPX, hydrogen peroxide; TBH, tert-butyl hydroperoxide; DIA, diamide; MD, menadione; MMS, methyl methanesulfonate; HU, hydroxyurea; TM, tunicamycin; DTT, dithiothreitol; CR, Congo red; CFW, calcofluor white; SDS, sodium dodecyl sulfate; CDS, cadmium sulfate; KCR, YPD + KCl; NCR, YPD + NaCl; SBR, YPD + sorbitol; KCS, YP + KCl; NCS, YP + NaCl; SBS, YP + sorbitol. Red and blue gradients represent phenotype reduction and enhancement, respectively, with strong, intermediate and weak phenotypes indicated by color intensity. (D) Growth in Sabouraud Dextrose (SabDex) medium showing cell aggregation in WT and *cna1*Δ strain across different clades.

To validate the functions of Cna1 in clades other than clade I, we constructed *CNA1* knockout mutants in strains B11220 (Clade II), B11221 (Clade III), and B11245 (Clade IV) (S6 Fig). Spot assays were conducted to evaluate the mutants’ responses to various stress conditions. The stress response phenotypes of these mutants were largely consistent with those of B8441 (clade I), except for differences observed in their sensitivity to the cell wall-destabilizing agent, calcofluor white (CFW) (Fig 3B and C). Unlike *cna1*Δ mutants in clades I and II, *cna1*Δ mutants in clades III and IV exhibited increased resistance to CFW, implying that the calcineurin pathway may have clade-specific roles in maintaining cell wall integrity in *C. auris*.

Cell aggregation is a clade-specific phenotypic trait in *C. auris*, particularly prominent in clade III, where it is recognized as a key feature influencing virulence, antifungal susceptibility, and biofilm formation (18). To explore the role of calcineurin in this trait, we examined the aggregation ability of the *cna1*Δ mutants across different clades. Deletion of *CNA1* resulted in a general increase in cellular aggregation (Fig 3D). Notably, the clade III *cna1*Δ mutant exhibited a striking enhancement in aggregation compared to the wild-type strain (Fig 3D), which might contribute to its increased resistance to CFW (Fig. 3C). All these findings demonstrate that the calcineurin pathway plays both evolutionally conserved and clade-specific divergent roles in *C. auris*.

### Crz1 is the transcription factor downstream of calcineurin in *C. auris*

To identify transcription factors downstream of calcineurin, we queried the *Candida* genome database and discovered two potential Crz1 orthologs in *C. auris*: B9J08_002096 and B9J08_000447. B9J08_002096 was designated as Crz1 because it shares a closer phylogenetic relationship with *C. albicans* Crz1 than B9J08_000447, which we named Crz2 (S7A Fig). Both Crz1 and Crz2 contain conserved zinc finger C_2_H_2_ domains, which are essential for DNA binding (S7B, S7C, and S7D Fig). To elucidate the roles of Crz1 and Crz2 in *C. auris*, we generated *crz1*Δ, *crz2*Δ, and *crz1*Δ *crz2*Δ mutants using the B8441 strain as the parental strain (S8A, S8B, and S8C Fig).

The *crz1*Δ mutant displayed moderate sensitivity to fludioxonil and ER stress inducers, such as tunicamycin and DTT, compared to the calcineurin mutant (Fig 4A). Additionally, the *crz1*Δ mutant exhibited highly increased sensitivity to cell membrane stress induced by SDS, comparable to that observed in the *cnb1*Δ *cna1*Δ mutant (Fig 4A). Moreover, the *crz1*Δ mutant was more susceptible to azole drugs, including FLC, PSC, VRC, and KTC, than the wild-type strain, exhibiting phenotypes similar to the *cnb1*Δ *cna1*Δ mutant but with reduced severity (Fig 4A). Complementation with the wild-type *CZR1* allele restored normal phenotypes in the *crz1*Δ mutant (Fig 4A and S8D Fig). In contrast, the *crz2*Δ mutant did not exhibit any phenotypic alterations (Fig 4A). Furthermore, the *crz1*Δ *crz2*Δ double mutant exhibited phenotypes identical to those of the *crz1*Δ mutant alone. All these data suggest that Crz1, but not Crz2, likely functions as the downstream effector of calcineurin in *C. auris*.

**Fig 4.**
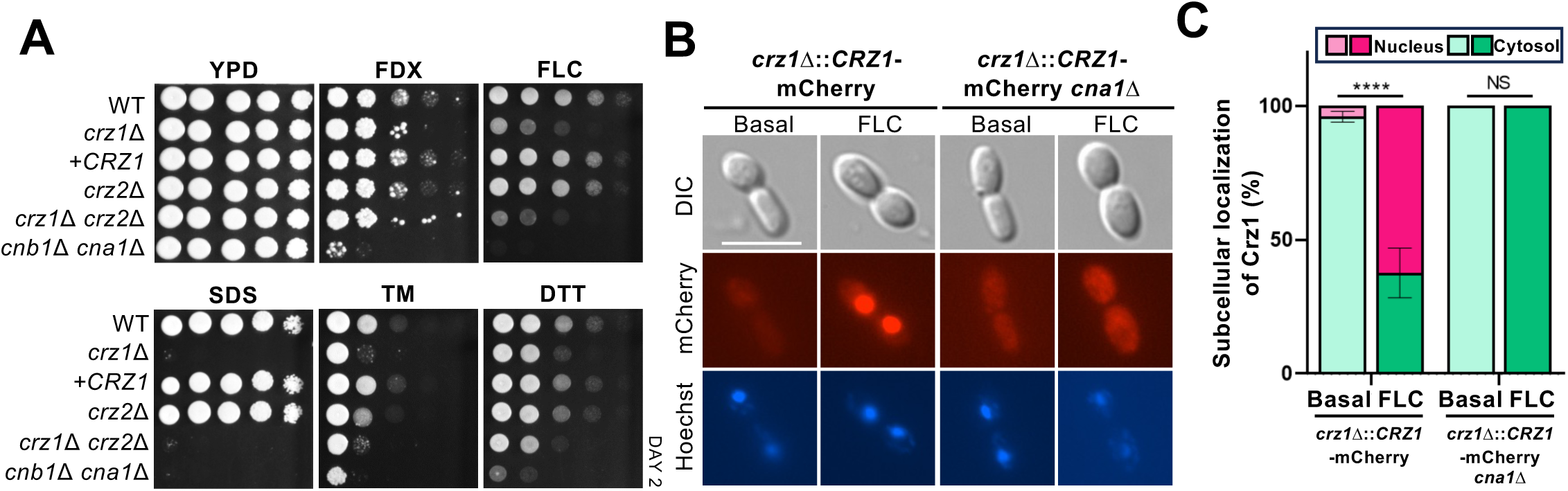
Crz1 functions as the downstream transcription factor of calcineurin in *C. auris*. (A) Qualitative spot assays showing the stress susceptibility of WT (B8441), *crz1*Δ (YSBA105), *crz1*Δ::*CRZ1* (YSBA158), *crz2*Δ (YSBA143), *crz1*Δ *crz2*Δ (YSBA153), and *cnb1*Δ *cna1*Δ (YSBA172) strains. WT and mutant strains were grown overnight in liquid YPD medium at 30°C, serially diluted 10-fold, and spotted on YPD medium containing stressors, including 3 μg/ml FDX, 150 μg/ml FLC, 0.1% SDS, 3.6 μg/ml TM, and 22 mM DTT. Plates were incubated for 2 days at 30°C. (B) Nuclear translocation of Crz1-mCherry upon FLC treatment. Cells were cultured overnight, synchronized to an OD_600_ of 0.2, and further grown until an OD_600_ of 0.8 was reached. Cells were treated with 400 μg/ml FLC for 3 h, harvested (1 ml), fixed, stained with Hoechst, and imaged using a fluorescence microscope. The white scale bar represents 10 μm. (C) Bar chart quantifying the proportion of cells exhibiting Crz1-mCherry localization in either the nucleus or cytoplasm. A total of 50 cells were counted (n=50). Statistical significance was evaluated using one-way ANOVA Analysis. Data are presented as mean ± SEM (NS, no significant; ****, *P* < 0.0001).

In several other fungi, activated calcineurin dephosphorylates Crz1, facilitating its translocation into the nucleus (5). To investigate the cellular localization of Crz1 in *C. auris*, we constructed the *crz1*Δ::*CRZ1-mCherry* strain, fin which the *crz1*Δ mutant was complemented with a C-terminal mCherry-tagged *CRZ1* allele (S9A Fig). The *CRZ1-mCherry* allele was functional, as it restored wild-type phenotypes in the *crz1*Δ mutant (S9C Fig). Upon FLC treatment for up to 4 h, the Crz1-mCherry protein predominantly localized to the nucleus after 3 h (Fig 4B and C). To confirm that calcineurin mediates Crz1 nuclear translocation, we deleted *CNA1* in the Crz1-mCherry tagged complemented strain (S9B and S9C Fig). In the absence of *CNA1*, Crz1 failed to localize to the nucleus in response to FLC (Fig 4B and C). These findings demonstrate that Crz1 functions as the downstream transcription factor of calcineurin and undergoes nuclear translocation in response to azoles in a calcineurin-dependent manner.

### Crz1 plays distinct roles with calcineurin in resistance to cell wall-damaging stressors and echinocandins in *C. auris*

We identified an unexpected phenotypic difference between the *crz1*Δ and calcineurin mutants. The *crz1*Δ mutant exhibited markedly increased resistance to cell wall-damaging stressor CFW and echinocandins, including CAF, MIF, and ANF, which was in stark contrast to the heightened susceptibility observed in the *cna1*Δ and *cnb1*Δ mutants (Fig 5A). Interestingly, Crz1 was still able to undergo nuclear translocation in response to CAF in a calcineurin-dependent manner (Fig 5B and C). These seemingly contradictory findings suggest that Crz1 may act as both a transcriptional activator and repressor, depending on the external cues and environmental context. Alternatively, Crz1 may regulate other transcription factors that, in turn, modulate genes involved in maintaining cell wall integrity in *C. auris*.

**Fig 5.**
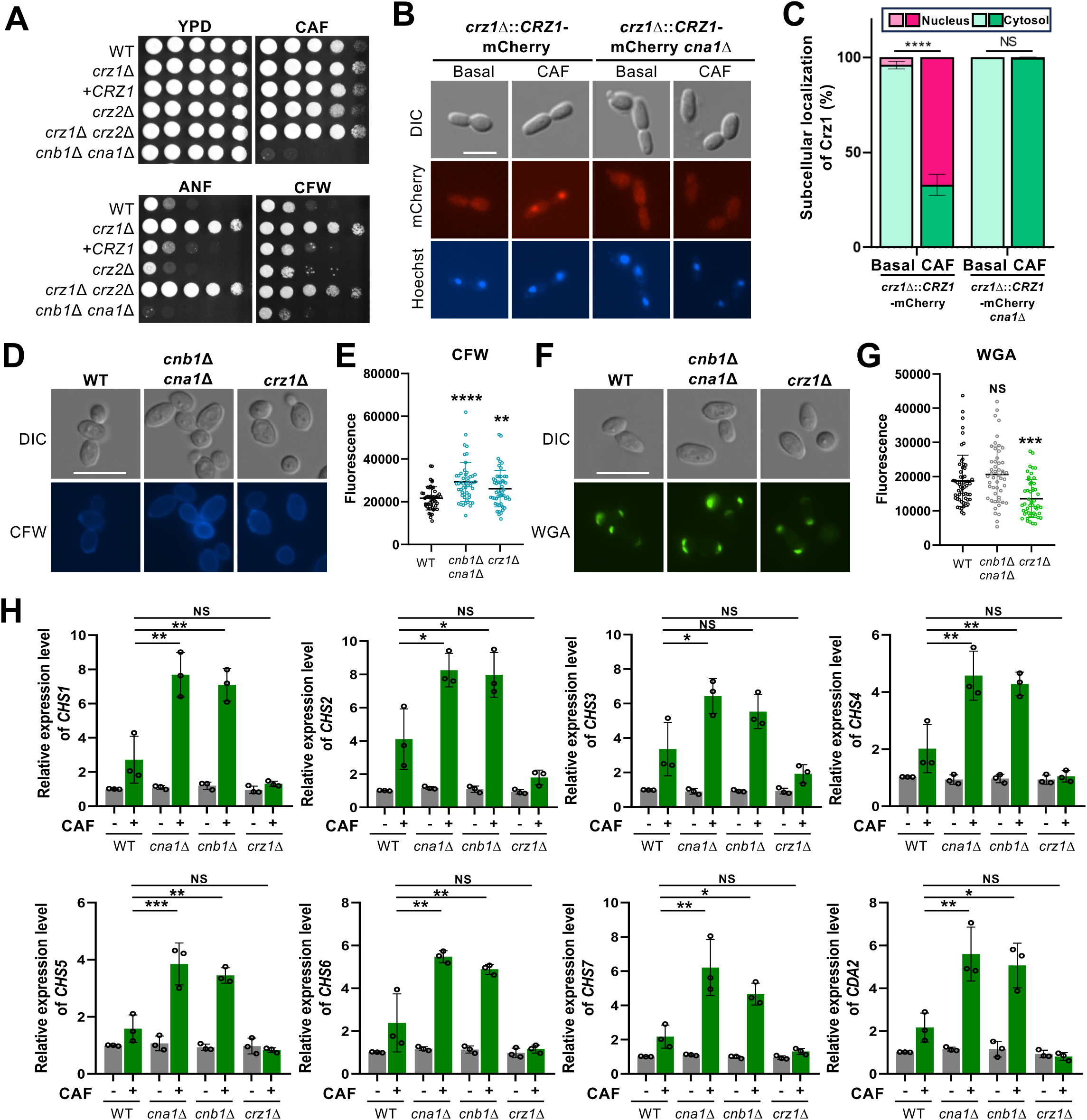
Crz1 opposes calcineurin in regulating echinocandin resistance in *C. auris*. (A) Spot assays showing the growth of WT and mutant strains on YPD medium containing 2.5 μg/ml CAF, 1 μg/ml ANF, or 0.12 mg/ml CFW. Plates were incubated for 2 days at 30°C. (B) Nuclear translocation of Crz1-mCherry upon treatment with 10 μg/ml CAF for 30 min. (C) Bar chart quantifying the proportion of cells exhibiting Crz1-mCherry localization in either the nucleus or cytoplasm. A total of 50 cells were analyzed (*n* = 50). Statistical significance was determined using one-way ANOVA. Data are shown as mean ± SEM (NS, no significant; ****, *P* < 0.0001). (D, F) Chitin and chito-oligomers staining in the WT and mutant strains. Cells were grown overnight at 30°C in liquid YPD medium, subcultured to an OD_600_ of 0.8, and stained with wheat germ agglutinin (WGA) or CFW. Representative fluorescence microscopy images are displayed for each strain (scale bar, 10 μm). (E, G) Quantitative fluorescence measurements of at least 50 individual cells per strain, analyzed using ImageJ/Fiji software, are shown. (H) qRT-PCR analysis of chitin synthase genes (*CHS1-CHS7*) and chitin deacetylase gene (*CDA2*) in WT, *cna1*Δ, *cnb1*Δ, and *crz1*Δ strains. Cells were cultured for 24 h with 5 μg/ml CAF at 30°C in a shaking incubator. Gene expression levels were normalized to *ACT1*, and fold changes were calculated relative to the basal expression level in WT. (E, G, H) Statistical significance was evaluated using one-way ANOVA with Bonferroni’s multiple-comparison test (*, *P* < 0.05; **, *P* < 0.01; ***, *P* < 0.001; ****, *P* < 0.0001; NS, not significant).

These findings suggest that calcineurin and Crz1 may play distinct roles in maintaining fungal cell wall integrity, as CFW and echinocandins disrupt chitin and β-glucan layers, respectively, which are major components of the fungal cell wall (19). To further investigate the differential roles of calcineurin and Crz1 in maintaining cell wall integrity, we analyzed CFW and wheat germ agglutinin (WGA) staining patterns in the wild-type, calcineurin (*cna1*Δ *cnb1*Δ), and *crz1*Δ mutant strains. CFW binds to β-linked polysaccharides, such as chitin and chitosan, while WGA with high molecular weight specifically binds to surface-exposed chito-oligomers (20, 21). CFW staining revealed increased fluorescence in both calcineurin and *crz1*Δ mutants compared to the wild type, indicating enhanced chitin accumulation in these mutants (Fig 5D and 5E). However, WGA staining showed that fluorescence levels in the calcineurin mutants were similar to those of the wild type, whereas the *crz1*Δ mutant exhibited significantly reduced fluorescence (Fig 5F and G). These results suggest that deletion of *CRZ1*, but not calcineurin, reduces chito-oligomer exposure.

To further investigate the roles of calcineurin and Crz1 in chitin biosynthesis, we measured the expression levels of the chitin synthase (*CHS*) and chitin deacetylase (*CDA*) genes. Under basal conditions, no significant changes in expression levels were observed (Fig 5H). However, upon treatment with CAF, the expression levels of chitin synthase genes (*CHS1*-*CHS7*) and *CDA2* were significantly upregulated in the *cna1*Δ or *cnb1*Δ mutants compared to the wild-type strain, whereas the *crz1*Δ mutant showed no notable increase (Fig 5H). These findings suggest that calcineurin mutants compensate for this heightened echinocandin susceptibility by upregulating chitin synthesis genes to reinforce the cell wall. In contrast, the *crz1*Δ mutant, which exhibits resistance to echinocandins, did not induce chitin synthesis under these conditions (Fig 5H). These findings underscore the complexity of the calcineurin/Crz1 pathway in maintaining fungal cell wall integrity.

### The calcineurin pathway is dispensable for morphogenesis, biofilm formation, secreted aspartyl protease activity, and ploidy switching in *C. auris*

We next examined whether the calcineurin/Crz1 pathway influences other known virulence traits in *C. auris.* Morphological changes, a critical virulence factors in *Candida* species, were assessed by inducing pseudohyphal formation with the DNA-damaging agent hydroxyurea (22). Both calcineurin and *crz1*Δ mutants exhibited no significant differences in pseudohyphal formation compared to the wild type (S10A Fig). Biofilm formation and the production of secreted aspartyl proteases (SAPs) were also analyzed in these mutants. Our findings confirmed that the calcineurin/Crz1 pathway does not play a role in either process (S10B and S10C Fig).

In addition, we investigated the frequency of ploidy switching, another virulence trait in *C. auris* (23). It has been reported that haploid and diploid *C. auris* cells grown on YPD medium containing phloxine B display white and pink colony phenotypes, respectively. Interestingly, most calcineurin mutants, but not the *crz1*Δ *mutant*, formed pink colonies (S10D Fig). However, fluorescence-activated cell sorting (FACS) analysis revealed that the pink-colored calcineurin mutant cells were not diploid (Fig S10D Fig), suggesting that ploidy switching did not occur in these strains. We hypothesized that the pink colony phenotype may result from altered cell membrane or wall composition, which allows the dye to penetrate more easily in the calcineurin mutant strains compared to the wild-type strain. Collectively, these results demonstrate that the calcineurin/Crz1 pathway is not involved in biofilm formation, secreted aspartyl protease activity, and ploidy switching in *C. auris*.

### Roles of the calcineurin pathway in the pathogenicity of *C. auris*

To determine the role of the calcineurin pathway in the pathogenicity of *C. auris*, we first utilized the *Drosophila melanogaster* infection model, a well-established fungal infection system, recognized for its short lifecycle, cost-effectiveness, ease of handling, and genetic consistency (24). A total of 80 flies per group were injected with wild-type, *cna1*Δ, *cna1*Δ::*CNA1*, or *byc1*Δ strains, and the survival rates were monitored over four days (Fig 6A). The *bcy1*Δ (YSBA4) strain, which lacks the regulatory subunit of protein kinase A (PKA) in the cAMP pathway, was included as a control strain due to its previously reported attenuated virulence in murine systemic infection models (25). Flies infected with the *cna1*Δ mutants exhibited a significantly higher survival rate than those infected with the wild-type and *cna1*Δ::*CNA1* strains. However, this increased survival was less pronounced than in flies infected with the *bcy1*Δ control strain (Fig 6B). These results indicate that the calcineurin pathway is required for the virulence of *C. auris* in the insect infection model.

**Fig 6.**
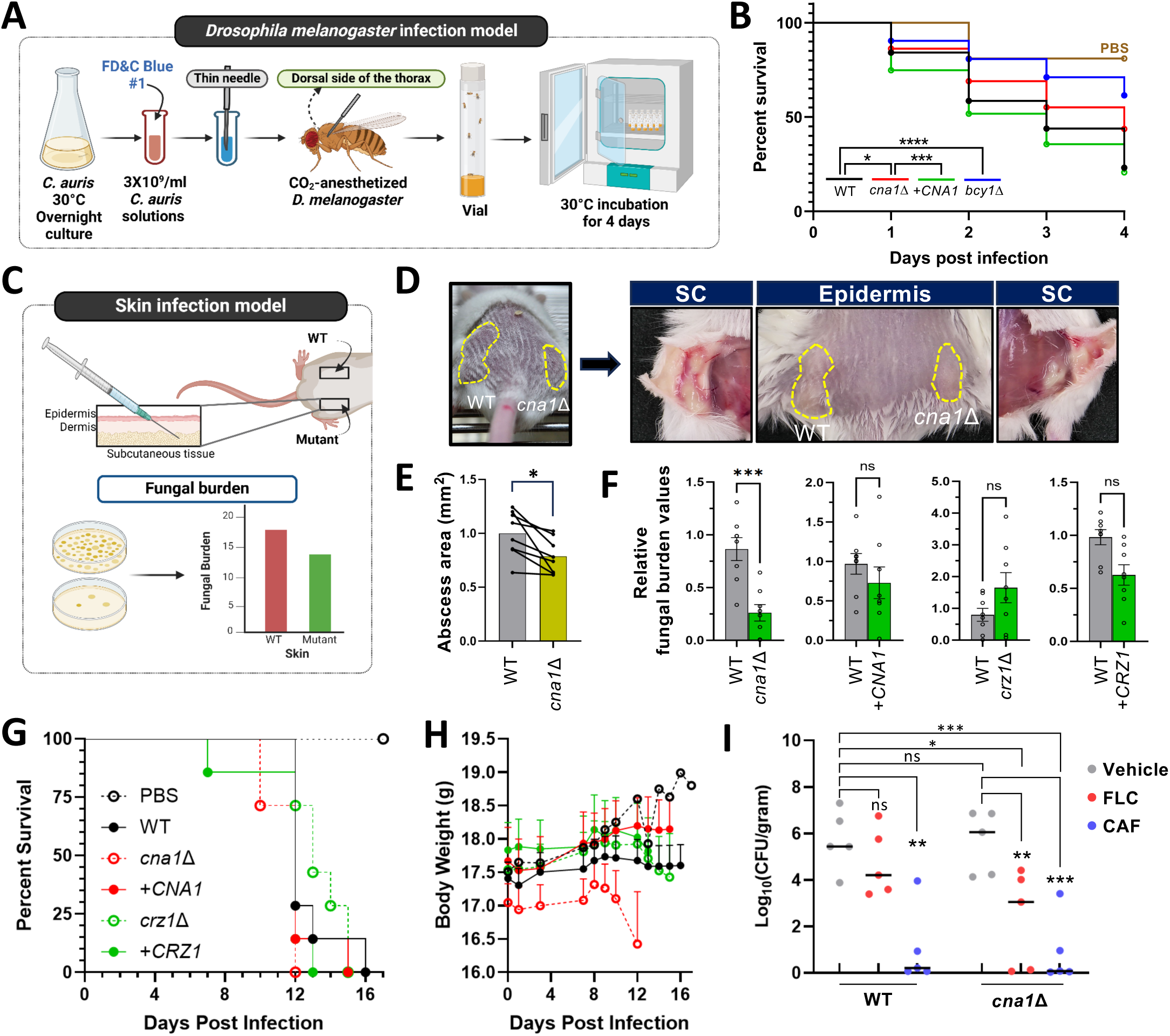
The role of the calcineurin pathway in the pathogenicity of *C. auris*. (A) Schematic representation of the experiment processes for the *Drosophila melanogaster* infection model. (B) Survival rates of *D. melanogaster* infected with WT, *cna1*Δ, +*CNA1*, or *bcy1*Δ strains were monitored over 4 days post-infection (dpi). A total of 80 files were used per strain. Survival curves were statistically analyzed using the log-rank (Mantel-Cox) test. The PBS control is represented in brown. (C) Schematic diagram of the mouse skin infection model. (D) A total of 10^7^ cells from WT or mutant strains were subcutaneously injected into the left and right flanks of 7-week-old BALB/c mice (n=8). (E) Clinical scores were determined by measuring abscess areas. Statistical significance was calculated using an unpaired *t*-test (*, *P*=0.0424). (F) Fungal burden was assessed by homogenizing abscess tissue 9 dpi and plating the homogenate on YPD agar plate. Relative fungal burden values were calculated as mutant CFU/g per WT CFU/g. Statistical significance was calculated using an unpaired *t*-test (***, *P* < 0.0005). (G, H) Survival curve and body weight changes of 7-week-old BALB/c mice (n=7) intravenously injected with 10^7^ *C. auris* cells. (I) Fungal burden in the inner ear was measured 7 days post-intravenous systemic infection. Mice were treated with FLC (20 mg/kg) and CAF (2 mg/kg) via intraperitoneal injection, administered once daily for six days, starting on the day of infection. Statistical significance was calculated using an unpaired *t*-test (**, *P* < 0.01; ***, *P* < 0.001)

We next evaluated the virulence potential of the calcineurin mutants using a murine (BALB/c) infection model. The skin is the primary colonizing niche for *C. auris*. Given that calcineurin mutants exhibit defects in maintaining cell wall integrity, which may impair fungal colonization of the skin, we employed a subcutaneous murine infection model using wild-type, *cna1*Δ, *cna1*Δ::*CNA1*, *crz1*Δ, or *crz1*Δ::*CRZ1* strains (Fig 6C). Fungal burdens in infected skin regions were quantified 9 days post-infection (dpi). Mice infected with the *cna1*Δ mutant exhibited significantly reduced abscess size and fungal burden compared to those infected with the wild-type strain. In contrast, mice infected with the *crz1*Δ mutant showed no significant differences in fungal burden relative to the wild type (Fig 6D-F). These results highlight the critical role of Cna1 as a virulence factor necessary for *C. auris* skin colonization, whereas Crz1 appears dispensable in this context.

Notably, the calcineurin mutant displayed a distinct virulence pattern in a systemic murine infection model compared to the skin infection model. Mice intravenously infected with the *cna1*Δ mutant exhibited survival rates similar to those infected with the wild-type or *cna1*Δ::*CNA1* strain (Fig 6G). However, noticeable body weight loss in *cna1*Δ-infected mice became apparent after 8 dpi, in contrast to mice infected with the wild-type or *cna1*Δ::*CNA1* strains (Fig 6H). In a systemic infection model treated with FLC and CAF, the fungal burden of the *cna1*Δ mutant in the inner ear was significantly more reduced by FLC compared to the wild-type strain (Fig 6I). Collectively, these findings suggest that the calcineurin pathway plays distinct roles in the virulence of *C. auris*, depending on the specific host niche.

## Discussion

This study investigates the critical role of the calcineurin signaling pathway in *C. auris*, emphasizing its essential functions in maintaining cell membrane and cell wall integrity, mediating antifungal resistance, facilitating environmental stress adaptation, and contributing to pathogenicity (summarized in Figure 7). Calcineurin deletion (*cna1*Δ and *cnb1*Δ) results in pronounced hypersensitivity to high temperatures (>43°C) and cell membrane- and cell wall-disrupting agents. Loss of calcineurin markedly increases susceptibility to azoles and echinocandins, while enhancing resistance to amphotericin B, underscoring its potential as a therapeutic target. This is further reinforced by the synergistic antifungal effects observed when calcineurin inhibitors, such as FK506 or cyclosporin A, are combined with echinocandins or azoles. Crz1, the downstream transcription factor of calcineurin in *C. auris*, plays a central role in mediating responses to membrane and ER stress and in modulating drug susceptibility. Deletion of *CRZ1* increases susceptibility to azoles and membrane stressors but enhances resistance to echinocandins, suggesting that Crz1’s functions oppose certain aspects of calcineurin’s role. While calcineurin is dispensable for systemic infection in a mammalian host, it plays an important role in skin colonization, further supporting its potential as a universal antifungal target against multidrug-resistant fungal pathogens.

**Fig 7.**
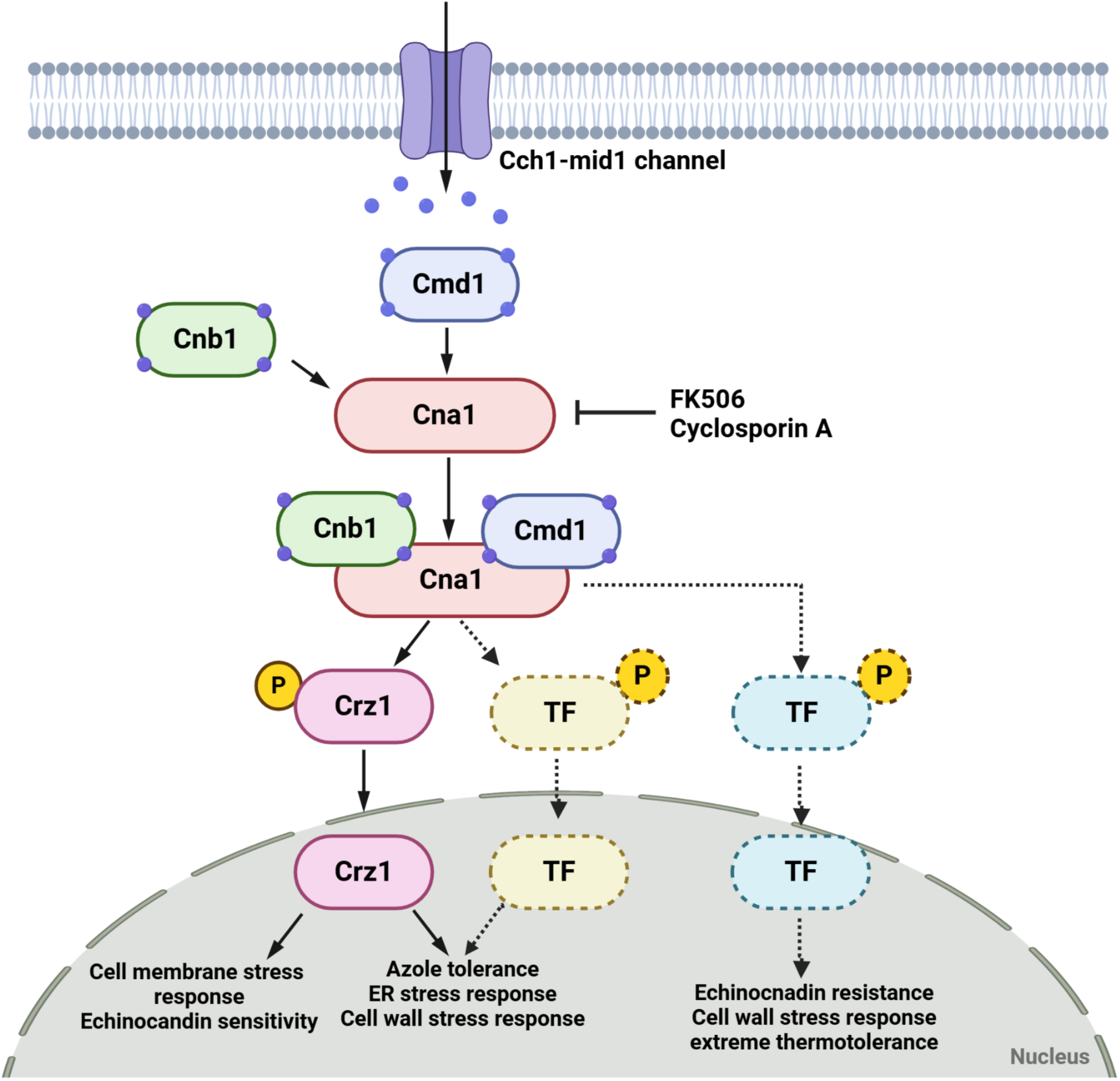
Proposed regulatory mechanisms of the calcineurin pathway in *C. auris*. The calcineurin pathway, consisting of the catalytic subunit Cna1 and the regulatory subunit Cnb1, regulates cell membrane and wall integrity, antifungal resistance, and virulence in *C. auris*. Upon calcium influx, activated calcineurin dephosphorylates Crz1, which translocates to the nucleus to modulate gene expression. While the calcineurin pathway promotes azole resistance, it exhibits antagonistic interactions with Crz1 in the regulation of echinocandin resistance and cell wall integrity maintenance. These findings underscore both the conserved and unique regulatory roles of the calcineurin pathway in *C. auris*.

In *C. auris*, the calcineurin pathway shares several conserved roles with other *Candida* species (*C. albicans*, *C. glabrata*, *Candida tropicalis*, and *Candida dubliniensis*) as well as major fungal pathogens such as *Aspergillus fumigatus* and *Cryptococcus neoformans*. These roles include maintaining cell membrane and wall integrity, mediating resistance to environmental stresses, and promoting antifungal drug resistance, particularly to azoles and echinocandins. However, *C. auris* exhibits unique species-specific characteristics. For instance, in *C. neoformans*, calcineurin controls key developmental processes essential for their pathogenicity (9). In *A. fumigatus*, calcineurin controls normal filamentous growth (26). Furthermore, in *C. dubliniensis* and *C. tropicalis*, calcineurin plays a crucial role in hyphal growth (27). In contrast, our findings reveal that the calcineurin pathway in *C. auris* does not regulate morphogenic transitions. These results highlight both the conserved and distinct roles of the calcineurin pathway in *C. auris*, underscoring the pathogen’s specialized adaptation to its unique niche and stress environments.

Another unique feature of the *C. auris* calcineurin pathway is the regulatory mechanism of its downstream transcription factor Crz1. In *C. auris*, Crz1 functions downstream of calcineurin to mediate stress and drug responses through nuclear translocation. Crz1 plays a critical role in maintaining membrane integrity and regulating azole susceptibility, as its deletion phenocopies the effects observed in calcineurin mutants. Similarly, in *C. neoformans*, calcineurin regulates stress survival and virulence through Crz1, along with additional downstream targets, including Lhp1, Pbp1, and Puf4, which contribute to post-transcriptional regulation and stress adaptation (9). However, a distinct feature in *C. auris* is the opposing effects of calcineurin and Crz1 on echinocandin resistance: calcineurin deletion increases echinocandin susceptibility, whereas Crz1 deletion enhances resistance. This phenotype is highly unusual and contrasts with other fungal species, such as *C. albicans* and *C. glabrata*, where both calcineurin and Crz1 are required for echinocandin resistance (8, 28). These findings suggest that Crz1 may function as both an activator and a repressor, regulating distinct gene sets that influence echinocandin resistance. Alternatively, calcineurin may activate additional transcription factor(s) to promote echinocandin resistance. Although we identified Crz2, a structural paralog of Crz1, its deletion did not affect any calcineurin-dependent phenotypes. Further investigation is warranted to identify additional calcineurin-dependent transcription factors beyond Crz1 in *C. auris*.

Our findings further demonstrate that calcineurin and Crz1 play distinct roles in regulating cell wall composition in *C. auris*. Both the *cnb1*Δ *cna1*Δ and *crz1*Δ mutants exhibited increased CFW staining, indicating elevated accumulation of chitin and its derivative. However, the *crz1*Δ mutants showed reduced WGA fluorescence staining compared to the wild-type and calcineurin mutants. Given that WGA is a high molecular weight lectin that specifically stains surface-exposed chitins, such as chito-oligomers, deletion of *CRZ1* deletion may result in diminished surface exposure of chito-oligomers. This reduction in surface-exposed chito-oligomers could be attributed to the increased beta-glucan thickness observed in the *crz1*Δ mutants, which may also explain the enhanced echinocandin resistance in these mutants. These observations indicate that Crz1-mediated regulation of cell wall composition likely influences both chitin exposure and echinocandin resistance. Further investigations are required to elucidate differential roles of calcineurin and Crz1 in modulating cell wall architecture and composition in *C. auris*.

The conservation of calcineurin functions across *C. auris* clades highlights its critical role in stress adaptation and survival in this fungal pathogen. Phylogenetic analysis confirmed that Cna1 is highly conserved, while functional assays demonstrated consistent stress response and antifungal drug resistance phenotypes across clades. These findings underscore calcineurin as an evolutionarily conserved regulator and a promising antifungal target effective across the genetically diverse *C. auris* clades. Interestingly, a clade-specific variation in calcineurin function was observed in sensitivity to CFW. While clades I and II displayed enhanced sensitivity, clades III and IV showed reduced sensitivity. Since CFW is a cell wall-destabilizing agent that binds to chitin and its derivative, these results suggest that the calcineurin pathway may have differential roles in maintaining cell wall integrity across clades. This variation may correlate with cell aggregation phenotype predominantly observed in clade III strains (20), implying potential differences in cell surface structures among *C. auris* clades. Further investigation into calcineurin-dependent clade-specific traits is warranted to understand the underlying mechanisms driving these phenotypic differences.

We observed a notable difference in the virulence potential of calcineurin across varying animal model systems. The *cna1*Δ mutant exhibited reduced virulence in both the *Drosophila* infection model and the murine skin infection model. In contrast, the *cna1*Δ mutant retained virulence comparable to the wild-type strain in a systemic murine infection model. Interestingly, mice intravenously infected with the *cna1*Δ mutant experienced significant body weight loss during infection, a phenomenon not observed in mice infected with the wild-type or its complemented strains. As Crz1 was dispensable for virulence in all tested animal models, we hypothesized that the opposing roles of calcineurin and Crz1 in maintaining cell wall integrity could influence virulence outcomes. This finding contrasts with the established role of calcineurin in systemic infections caused by other fungal pathogens, such as *C. albicans* and *C. neoformans*, where it is essential for adaptation to host environments and virulence (29, 30). We speculate that the altered cell wall integrity in the *cna1*Δ mutant may provoke an abnormal immune response during systemic infection of *C. auris*, contributing to a significant body weight loss. A similar immune response phenotype has been reported in *rim101* mutants of *C. neoformans* (31). Further studies are required to elucidate how altered calcineurin signaling in *C. auris* impacts host immune systems during systemic infection.

Based on our data, calcineurin inhibitors may not be useful for treating systemic infection caused by *C. auris* but could prove effective for managing its skin infections or other superficial or cutaneous infections. Our data demonstrated that calcineurin inhibitors, such as FK506 and cyclosporin A, synergize with azoles and echinocandins to confer fungicidal activity against *C. auris*. Notably, we observed a significant reduction in the fungal burden of the *cna1*Δ mutant in the inner ear by FLC treatment, suggesting that the combination of azoles and calcineurin inhibitors could exhibit synergistic effects in vivo for treating *C. auris* infections in the inner ear. In *C. albicans*, FK506 and cyclosporin A improve FLC efficacy by impairing stress adaptation mechanisms (32, 33). Similarly, in *C. neoformans* and *A. fumigatus*, FK506 enhances azole and echinocandin activity by disrupting stress tolerance and cell wall remodeling (34, 35). In *C. glabrata*, FK506 synergizes with FLC by impairing stress adaptation (36). Despite its potent antifungal activity, FK506 is limited by its immunosuppressive effects, prompting the development of analogs designed to reduce immunosuppression while selectively targeting fungal cells (37, 38). These FK506 analogs have demonstrated synergistic effects with FLC in *C. albicans* and *C. neoformans* (38). While FK506’s immunosuppressive properties remain a challenge, our findings underscore the potential of calcineurin inhibitors as adjunctive agents to enhance the efficacy of azoles and echinocandins for *C. auris* infections, particularly in non-systemic settings.

In conclusion, our study demonstrates that the calcineurin complex and its downstream transcription factor Crz1 play overlapping and distinct roles in regulating extreme thermotolerance, cell membrane and wall integrity, antifungal drug resistance, and virulence in *C. auris*. Further genetic, transcriptomic, and biochemical investigations are warranted to elucidate the precise mechanisms underlying the calcineurin complex and its downstream regulatory network in *C. auris*.

## Materials and Methods

### Ethics statement

All animal care and experimental procedures were reviewed and approved by the Institutional Animal Care and Use Committee of the Experimental Animal Center at Jeonbuk National University (Approval number JBNU 2023–131). All experiments were conducted in full compliance with established ethical guidelines for animal research.

### *Candida auris* strains and growth media

The *Candida auris* strains used in this research are listed in S1 Table in the supplemental material. The parental wild-type strains, including B8441 (clade I, AR0387), B11220 (clade II, AR0381), B11221 (clade III, AR0383), and B11245 (clade IV, AR0386), were provided by the Centers for Disease Control and Prevention (CDC). Both wild-type and mutant strains were maintained as frozen stocks in 20% glycerol at −80°C until use. Yeast strains were cultured on YPD agar plates (2% agar in YPD broth: 2% peptone, 1% yeast extract, and 2% D-glucose) at 30°C for 24 to 48 h. For liquid culture preparation, yeast cells were grown in YPD media at 30°C with constant agitation at 200 rpm. Before experimental assays, cells were transferred to fresh YPD broth and cultured to the mid-log phase (OD_600_ = 0.6–0.8) prior to the designated treatments.

### Gene deletion and complementation

Gene deletion mutants were constructed using the nourseothricin resistance (*CaNAT*), hygromycin B resistance (*CaHYG*), or G418 resistance (*CaNEO*) markers, each flanked by 0.5–0.7 kb 5′ and 3′ regions of the target genes, including *CNA1, CNB1, CRZ1,* and *CRZ2*. Gene disruption cassettes containing the selection markers were assembled through double-joint PCR. In the first round of PCR, the flanking regions of each target gene were amplified using the L1-L2 and R1-R2 primer pairs. The *CaNAT* selection marker was amplified from the plasmid pV1025, which contains the *CaNAT* gene, using the appropriate primer pairs listed in S1 Table in the supplemental material. Similarly, the *CaNEO* selection marker was amplified from the plasmid pTO149 RFP-NEO, which carries the *CaNEO* gene, using corresponding primer pairs. The PCR products of the flanking regions and selection markers were purified and subsequently used as templates for a second round of double-joint PCR. During this step, the 5′- and 3′-gene disruption cassettes containing split selection markers were amplified using the L1-split primer 2 and R2-split primer 1, respectively (see S1 Table in the supplemental material).

Gene disruption cassettes were introduced into *C. auris* using a modified lithium acetate/heat-shock transformation protocol. Overnight cultures were grown at 30°C in 50 ml of YPD broth with constant shaking. A 1.2 ml aliquot of cultured cells was harvested by centrifugation, washed sequentially with dH_2_O and lithium acetate buffer (100 mM lithium acetate, 10 mM Tris, 1 mM EDTA, pH 7.5), and resuspended in 300 μl of lithium acetate buffer. For each transformation, the reaction mixture consisted of 10 μl of denatured salmon sperm DNA (Sigma, cat no. D9156), 100 μl of the cells, 500 μl of 50% PEG4000 (Sigma, cat no. P4338), and 50 μl of the amplified gene deletion cassette. The mixture was incubated at 30°C for 6 h with intermittent vertexing, followed by a 20-min heat shock at 42°C and immediate cooling on ice for 1 min. The cells were then pelleted, resuspended in 1 ml of YPD medium, and incubated at 30°C for 1-2 h with shaking to allow recovery. After recovery, the cells were washed twice with fresh liquid YPD medium and plated onto selective YPD agar containing 400 μg/ml nourseothricin, 1.8 mg/ml hygromycin B, or 2.4 mg/ml G418. Plates were incubated at 37°C for 2–3 days to select transformants. Positive transformants resistant to nourseothricin, hygromycin B, or G418 were verified for the correct genotype using diagnostic PCR and confirmed via Southern blot analysis.

To validate the phenotypes of the *cna1*Δ, *cnb1*Δ, and *crz1*Δ mutants, we generated corresponding complemented strains by re-integrating the respective wild-type genes into their native loci (*cna1*Δ::*CNA1, cnb1*Δ::*CNB1*, and *crz1*Δ::*CRZ1*). Full-length gene fragments were amplified by Pfu-PCR using genomic DNA extracted from the wild-type B8441 strain as a template, along with the primer pairs listed in S2 Table in the supplemental material. The amplified fragments were cloned into the TOPO vector (Invitrogen), resulting in the plasmids pTOP-CNA1, pTOP-CNB1, and pTOP-CRZ1. After sequence verification, each plasmid was modified by subcloning the *CaHYG* selection marker into the corresponding pTOP vector, generating pTOP-CNA1-HYG, pTOP-CNB1-HYG, and pTOP-CRZ1-HYG. For re-integration into the native locus, pTOP-CNA1-HYG, pTOP-CNB1-HYG, and pTOP-CRZ1-HYG were linearized using PmlI, BsmI, and BglII, respectively, and introduced into the corresponding mutant strains using the lithium acetate heat-shock transformation protocol. The successful generation of complemented strains was confirmed through diagnostic PCR, verifying the correct integration of the wild-type alleles.

### Total RNA preparation and quantitative RT-PCR

Total RNA was extracted from *C. auris* wild-type and calcineurin/Crz1 mutant strains cultured overnight at 30°C in YPD broth. Cells were harvested by centrifugation at mid-log phase (OD_600_ of 0.6–0.8), rapidly frozen in liquid nitrogen, and lyophilized. For RNA isolation, 30 ml of the culture was maintained as the basal condition, while the remaining 30 ml was subjected to additional incubation with specific stress agents. Total RNA was extracted using the Trizol-based Easy Blue reagent (Intron). Complementary DNA (cDNA) was synthesized from purified total RNA using reverse transcriptase (Thermo Scientific) following the manufacturer’s protocol. Quantitative PCR (qPCR) was performed using a CFX96^TM^ Real-Time system (Bio-Rad) with gene-specific primer pairs. Expression levels were normalized to the housekeeping gene *ACT1*. Statistical analyses were conducted using one-way ANOVA followed by Bonferroni’s multiple-comparison test. To ensure reproducibility. All experiments were performed in technical triplicates and independently repeated three times with biological replicates.

### Growth and stress sensitivity spot assay

To evaluate the growth and stress sensitivity of wild-type and calcineurin/Crz1 mutant strains of *C. auris*, cells were cultured overnight at 30°C, serially diluted 10-fold up to four times (final dilution 1:10^4^), and spotted onto YPD plates. The plates were incubated at various temperatures (25°C, 30°C, 37°C, 42°C, and 45°C), and growth was qualitatively assessed by photographing the plates after 24 h of incubation. To assess stress responses, specific chemical stressors were incorporated into the growth media, including an osmotic stressor (sorbitol), cation and salt stressors (NaCl or KCl), oxidative stressors [hydrogen peroxide (HPX), *tert*-butyl hydroperoxide (TBH), menadione (MD), or diamide (DIA)], a membrane destabilizing stressor (SDS), cell-wall destabilizing stressors [Congo red (CR) and calcofluor white (CFW)], and various antifungal agents [fludioxonil (FDX), flucytosine (5FC), fluconazole (FLC), itraconazole (ITC), posaconazole (PSC), ketoconazole (KTC), itraconazole (ITC), caspofungin (CAF), micafungin (MIF), anidulafungin (ANF) or amphotericin B (AMB)]. The plates were grown at 30°C and photographed after 2-3 days of incubation under stress conditions.

### EUCAST MIC test and checkerboard assay

Wild-type and mutant *C. auris* strains were cultured overnight at 30°C in YPD medium, washed twice with dH_2_O, and resuspended in dH_2_O. For the EUCAST (European Committee on Antimicrobial Susceptibility Testing) MIC assay, the cell suspension was adjusted to an OD_600_ of 1.0. A total of 100 μl of the prepared cell suspension was added to 10 ml of 3-(*N*-morpholino)propanesulfonic acid (MOPS)-buffered RPMI 1640 media (pH 7.4, supplemented with 0.165 M MOPS and 2% glucose), and the mixture was dispensed into 96-well plates containing 2-fold serial dilution of the test drugs. For the checkerboard assay, the cell suspension in RPMI medium was dispensed into 96-well plates containing 2-fold serial concentrations of each drug combination. The plates were incubated at 35°C for 48 h. Cell density in each well was measured at OD_595_ to determine the MIC values. Following growth assessment, cultures from each well were spotted onto YPD agar plates and incubated at 30°C for 24 h to evaluate their fungicidal effects.

### Evaluating cell aggregation

Wild and mutant strains were cultured in Sabouraud Dextrose (SabDex) medium for two days, washed twice with phosphate-buffered saline (PBS), and counted. A total of 2.5 × 10^8^ cells were suspended in 5 ml PBS in sterile medical tubes, vortexed thoroughly, and photographed 5 min after vortexing (39). Images were captured using fluorescence microscopy.

### Observing the intracellular localization of mCherry-tagged proteins

Crz1-mCherry-tagged strains were grown overnight in YPD broth at 30°C. The overnight cultures were subcultured in YPD medium until an OD_600_ of 0.8 was reached. The cells were treated with antifungal drugs and incubated under the designated conditions. For cell fixation, the samples were treated with a 4% paraformaldehyde solution containing 3.4% sucrose for 15 min at room temperature. The fixed cells were thoroughly washed with a buffer containing 0.1 M KPO4 and 1.2 M sorbitol. The nucleus was visualized by staining the cells with 10 μg/ml Hoechst 33342 (Thermo Fisher, USA) for 30 min in the dark. Finally, the stained cells were examined using differential interference contrast (DIC) and fluorescence microscopy (Nikon Eclipse, Japan) to determine the subcellular localization of Crz1-mCherry fusion proteins.

### Measurement of chitin and chito-oligomers content in cell walls

To measure the chitin and chito-oligomers content in the cell walls of wild-type and calcineurin/Crz1 mutant strains of *C. auris*, cells were cultured overnight at 30°C in a shaking incubator. The cells were harvested by centrifugation, washed, and resuspended in PBS (pH 7.5). For staining, the cells were treated with 100 μg/ml fluorescein isothiocyanate (FITC)-conjugated WGA or 25 μg/ml CFW for 30 min in the dark. Following staining, cells were washed three times with PBS and visualized by fluorescence microscopy (Olympus BX51). Fluorescence intensity from at least 50 individual cells per sample was quantified using ImageJ/Fiji software to determine relative chitin and chito-oligomers levels.

### Assessment of morphological transition from yeasts to pseudohyphae

To induce pseudohyphal formation in wild-type and calcineurin/Crz1 mutant strains, cells were cultured for 24 h in YPD broth supplemented with 100 mM HU. Following pseudohyphal induction, the cells were fixed using 10% formalin and stained with 10 μg/ml Hoechst 33342 (Thermo Fisher) to visualize cellular morphology. The fixed and stained samples were incubated in the dark for 30 min, after which images were captured using fluorescence microscopy.

### Crystal violet assay for biofilm formation

*C. auris* wild-type and mutant strains were cultured overnight at 30°C in 2 ml of YPD liquid medium. The cells were washed twice with sterile H_2_O and resuspended in MOPS-buffered RPMI-1640 media (pH 7.4, 0.165 M MOPS and 2% glucose). The cell suspension was adjusted to an OD_600_ of 0.5, and 200 μl of the suspension was dispensed into each well of a 96-well plate. The cultures were incubated at 37°C with shaking at 220 rpm for 24 h. After incubation, the cell suspensions were carefully removed, and the plates were dried in a dry oven at 65°C. Each well was then treated with 150 μl of 0.5% crystal violet solution and incubated at room temperature for 10 min for staining. Excess dye was removed by washing the wells three times with PBS, and the plates were dried again in the dry oven at 65°C. To solubilize the bound crystal violet, 200 μl of 33% acetic acid was added to each well and incubated for 2 min. The solubilized solution was diluted 1:10 with PBS and transferred to a new 96-well plate. Absorbance was measured at OD_595_ using a microplate reader.

### Secreted aspartyl proteinase activity assay

SAP activity was assessed using the yeast carbon base-bovine serum albumin (YCB-BSA) method. *C. auris* strains were grown overnight in 2 ml of YPD broth at 30°C, harvested by centrifugation, washed with dH_2_O, and resuspended in 1 ml of dH_2_O. The cell suspension was adjusted to a final concentration of 10^5^ cells/ml. Subsequently, 3μl of the cell suspension was spotted onto YCB-BSA plates (containing 23.4 g/l yeast carbon base and 0.2% BSA) and incubated at various temperatures (30°C, 37°C, and 42°C) for 3 days. SAP activity was quantified by measuring the diameter of the halo formed around the colonies. All experiments were performed in triplicate with independent biological repeats to ensure reproducibility.

### Ploidy switching and flow cytometry analyses

*C. auris* wild-type and calcineurin/Crz1 mutant strains were cultured overnight at 30°C in YPD medium, washed twice with PBS, and counted. For the ploidy switching assay, 100 cells were spread onto YPD media supplemented with 5 μg/ml phloxine B and incubated at 25°C for 14 days. For DNA content analysis, single colonies from YPD medium containing phloxine B were inoculated into liquid YPD medium and cultured overnight at 30°C. The cells were then harvested, washed with PBS, and counted. A total of 10^6^ cells were fixed in 70% ethanol for 14-16 h at 4°C. Following fixation, cells were washed twice with PBS and treated with RNase A (200 μg/ml) at 37°C for 1 h to remove RNA contamination. After centrifugation, the supernatant was discarded, and the cells were stained with 500 μl of 100-μg/ml propidium iodide for 30 min at room temperature in the dark. The stained cells were washed with PBS, resuspended in 300 μl of PBS, and analyzed for DNA content using flow cytometry. A minimum of 10,000 cells were evaluated per experiment to ensure statistical reliability.

### Drosophila melanogaster infection model

The *Drosophila melanogaster* (w1118) was used for infection experiments. Flies were maintained at 25°C with a 12-h light-dark cycle on standard cornmeal-agar medium obtained from Bloomington Stock Center. Cells from *C. auris* wild-type and mutant strains were cultured overnight at 30°C in YPD medium, followed by three washes with PBS. The cell concentration was adjusted to 3 × 10^9^ cells/ml. The cell suspensions were centrifugated, and the resulting pellets were resuspended in PBS containing 1% (w/v) blue food dye (FD&C Blue #1) to ensure visualization. For the survival assay, adult female flies (4-5 days old) were anesthetized using CO_2_ and the dorsal thorax was pricked with a thin needle dipped in the prepared cell suspension. Approximately 80 flies per strain were infected, divided into groups of 10, and placed into individual vials containing standard cornmeal-agar medium. The flies were incubated at 30°C, and survival was monitored daily for up to 4 days post-infection (dpi).

### Subcutaneous murine infection models

*C. auris* wild-type and mutant strains were cultured overnight in YPD medium at 30°C with shaking. The cell concentration was adjusted to 10⁸ cells/ml, and 100 μl of the suspension in PBS was injected subcutaneously into anesthetized mice. SPF/VAF-confirmed, inbred 6-week-old female BALB/cAnNCrlOri mice (ORIENT BIO INC., South Korea) were housed for one week prior to infection. Anesthesia was induced by inhalation of isoflurane vapor (Hana Pharm. Co., Ltd., South Korea) at a flow rate of 80 cc/min. Subcutaneous injection was performed on both flanks of the mice, where the fur had been trimmed with clippers. The size of abscesses formed post-infection was quantified using ImageJ with a ruler as a reference. Colony-forming units (CFUs) were determined by surgically extracting the abscess, homogenizing the tissues, and plating the homogenates on YPD medium for colony enumeration.

### Systemic murine infection models

The cell concentration was adjusted to 10⁸ cells/ml, and 100 μl of the suspension in PBS was injected intravenously into restrained mice. Humane endpoints were defined by the onset of systemic infection symptoms, including rapid weight loss, abnormal head tilt, and circling behavior. Survival was monitored and represented as a survival curve. For drug treatment groups, drugs were administered intraperitoneally starting from the day of infection. Each drug was prepared in a 5% Kolliphor solution (Polyoxyl-35 castor oil:Ethanol = 1:1), and the injection volume was limited to 100 μl. The stock concentration for a 20 mg/kg (mpk) dose was 4 mg/ml, while for a 2 mpk dose, the stock concentration was 0.4 mg/ml.

## Acknowledgments

This work was supported by the National Research Foundation of Korea funded by the Korean government (MSIT) (2021R1A2B5B03086596 and 2021M3A9I4021434 to YSB; 2022R1C1C2003274 and 2021R1A6C101C369 to KTL; RS-2024-00345184 to WJL) and by the Yonsei Signature Research Cluster Program (2023-22-0012 to YSB). The funders had no role in study design, data collection and analysis, decision to publish, or preparation of the manuscript.

## Author contributions

### Hyunjin Cha

ROLES: Data curation, Methodology, Formal analysis, Investigation, Validation, Visualization, Writing – original draft

### Doyeon Won

ROLES: Data curation, Formal analysis, Investigation, Methodology

### Seun Kang

ROLES: Data curation, Formal analysis, Investigation, Methodology

### Eui-Seong Kim

ROLES: Data curation, Formal analysis, Investigation, Methodology

### Kyung-Ah Lee

ROLES: Data curation, Formal analysis, Investigation, Methodology

### Won-Jae Lee

ROLES: Data curation, Formal analysis, Funding acquisition, Methodology, Supervision, Writing – review & editing

### Kyung-Tae Lee

ROLES: Formal analysis, Funding acquisition, Investigation, Methodology, Resources, Supervision, Writing – review & editing

### Yong-Sun Bahn

ROLES: Conceptualization, Data curation, Formal analysis, Funding acquisition, Project administration, Resources, Supervision, Writing – original draft, review & editing

