## Supplemental Materials for "The calcineurin pathway regulates extreme thermotolerance, cell membrane and wall integrity, antifungal resistance, and virulence in *Candida auris*"

### S1 Table. Strains used in this study

| Strain | Genotype | Parent | Reference |
|---|---|---|---|
| **B8441** | Wild-type | | (1) |
| **B11220** | Wild-type | | (1) |
| **B11221** | Wild-type | | (1) |
| **B11245** | Wild-type | | (1) |
| **YSBA4** | *bcy1Δ::NAT* | B8441 | (2) |
| **YSBA24** | *tpk1Δ::NAT tpk2Δ::HYG* | YSBA17 | (2) |
| **YSBA119** | *sapa3Δ::NAT* | B8441 | (3) |
| **YSBA99** | *cna1Δ::NAT* | B8441 | This study |
| **YSBA102** | *cnb1Δ::NAT* | B8441 | This study |
| **YSBA105** | *crz1Δ::NAT* | B8441 | This study |
| **YSBA110** | *cna1Δ::CNA1_HYG* | YSBA99 | This study |
| **YSBA111** | *cnb1Δ::CNB1_HYG* | YSBA102 | This study |
| **YSBA143** | *crz2Δ::NAT* | B8441 | This study |
| **YSBA153** | *crz1Δ::NAT crz2Δ::NEO* | YSBA105 | This study |
| **YSBA158** | *crz1Δ::CRZ1_HYG* | YSBA105 | This study |
| **YSBA172** | *cnb1Δ::NAT cna1Δ::HYG* | YSBA102 | This study |
| **YSBA289** | *crz1Δ::CRZ1*-mCherry-*NEO* | YSBA105 | This study |
| **YSBA313** | *crz1Δ::CRZ1*-mCherry-*NEO cna1Δ* | YSBA289 | This study |
| **YSBA332** | *cna1Δ::NAT* | B11220 | This study |
| **YSBA336** | *cna1Δ::NAT* | B11221 | This study |
| **YSBA365** | *cna1Δ::NAT* | B11245 | This study |

### S2 Table. Primers used in this study

| Name | Primer description | Sequence (5’ to 3’) |
|---|---|---|
| **B11103** | pV1025 forward-extended primer | CTAGAACTAGTGGATCTGAA |
| **B11104** | pV1025 reverse-extended primer | TAAGAGTGAAATTCTGGAAA |
| **B11105** | NAT split primer 1 | CCATTGACTAAGGTTTTCCC |
| **B11106** | NAT split primer 2 | TTCAGTAGCCAAACCCATC |
| **B11107** | pV1025 Diagnostic screening primer 1 | TCAGTGGCAAATCCTAACC |
| **B11108** | pV1025 Diagnostic screening primer 1 | AGAGAAAATACCCGTGACG |
| **B12462** | pYM70 forward-extended primer 1 | CACATTTCCCCGAAAAGTGC |
| **B12463** | pYM70 reverse-extended primer 2 | TAACTTGCACTACCTCATCG |
| **B12464** | HYG split primer 1 | TGCTGATTTGTCTCAAACTT |
| **B12465** | HYG split primer 2 | TACCATTATCAGTCAAAACA |
| **B12486** | pYM70 Diagnostic screening primer 1 | TGCGGCACAATTGAATAGGG |
| **B12487** | pYM70 Diagnostic screening primer 1 | CGGTGATGACGGTGAAAACC |
| **B16376** | pTO149 forward-extended primer 1 | GACATGGAGGCCCAGAATAC |
| **B16377** | pTO149 reverse-extended primer 2 | CAGTATAGCGACCAGCATTC |
| **B16378** | NEO split primer 1 | TGCTCCTGCCGAGAAAGTAT |
| **B16379** | NEO split primer 2 | GCTCTTCGTCGAGATCATCC |
| **B16380** | pTO149 Diagnostic screening primer 1 | ACATGGGGATGTATGGGCTA |
| **B16381** | pTO149 Diagnostic screening primer 1 | TTTTCGCCTCGACATCATCT |
| **B14328** | *CNA1* 5’-flanking region primer L1 | TGTTGAAACACAAGGGCAAA |
| **B13428** | *CNA1* 5’-flanking region primer L2 | TTCAGATCCACTAGTTCTAGTTGTTATCGACGGGGGACAT |
| **B13429** | *CNA1* 3’-flanking region primer R1 | TTTCCAGAATTTCACTCTTATGAGAAAACTCTCTGGATAA |
| **B14329** | *CNA1* 3’-flanking region primer R2 | CCTCCCTTGGTCTTCTCACA |
| **B13431** | *CNA1* 5’-screening primer SO | GAACCTGGAAGATTCATGGT |
| **B13432** | *CNA1* 3’-sreening primer SO2 | TTTTCCTCTCAAAGCTTGCA |
| **B13433** | *CNA1* Southern blot probe primer PO | GCCTTCCTCTGTTACATAAA |
| **B13434** | *CNB1* 5’-flanking region primer L1 | GCTAGTCAAGAATGGCATCA |
| **B13435** | *CNB1* 5’-flanking region primer L2 | CAGATCCACTAGTTCTAGGTGGATGATGCTAACCCCAT |
| **B13436** | *CNB1* 3’-flanking region primer R1 | TCCAGAATTTCACTCTTATGACTTTGAACAATATTTAA |
| **B13437** | *CNB1* 3’-flanking region primer R2 | AGCAGTGTGCTTTCTTTACC |
| **B13438** | *CNB1* 5’-screening primer SO | CGACAGTGAACTCCTCGAAT |
| **B13439** | *CNB1* 3’-sreening primer SO2 | GACACCAAAGCTGTGATCAT |
| **B13440** | *CNB1* Southern blot probe primer PO | GGCCCCACTCCCATCGCTAT |
| **B13498** | *CRZ1* 5’-flanking region primer L1 | CCGGAGAGAAAATTGGATTT |

| | | |
|---|---|---|
| **B13499** | *CRZ1* 5'-flanking region primer L2 | TTCAGATCCACTAGTTCTAGTTCACGACGTTCAGAGACAT |
| **B13500** | *CRZ1* 3'-flanking region primer R1 | TTTCCAGAATTTCACTCTTACCATCTCGCCACCAGACTAG |
| **B13501** | *CRZ1* 3'-flanking region primer R2 | TCTTGTATTGATGAAAGCCA |
| **B13502** | *CRZ1* 5'-screening primer SO | GTGCATGAAAACACAGATCA |
| **B13503** | *CRZ1* 3'-sreening primer SO2 | TCTCCAATGCGATGTCGAAG |
| **B13504** | *CRZ1* Southern blot probe primer PO | GTCTTTGGAGAGGACCGAGG |
| **B16963** | *CNA1* Internal screening primer LP | AGCTATTGAGCCAGGAACCA |
| **B16964** | *CNA1* Internal screening primer RP | GGAGGGACCACGTAAAGACA |
| **B16574** | *CNB1* Internal screening primer LP | CCACGCTTCTAGAGTCCTTCA |
| **B16575** | *CNB1* Internal screening primer RP | TGTTCAAAGTCAACGCAGATG |
| **B16576** | *CRZ1* Internal screening primer LP | TCCCAACTCGTCATCCTACC |
| **B16577** | *CRZ1* Internal screening primer RP | CTGGCCAGGAGAATTCTGAG |
| **B16989** | CLP for *CNA1* complementation | CGATGGCGTGTATGGCAACA |
| **B16990** | CRP for *CNA1* complementation | GAAAGATGCGGTAGACCATA |
| **B16837** | CLP for *CNB1* complementation | GGTCTTTATGCTGCTGAAGG |
| **B16838** | CRP for *CNB1* complementation | AGTGGCTGAGAGGCAATTTG |
| **B16907** | CLP for *CRZ1* complementation | ATGAAAGCCGTTGTCATTGA |
| **B16908** | CRP for *CRZ1* complementation | CCAATGACATAACTTTTCCA |
| **B17049** | *CNA1* 5'-flanking region primer L1 (HYG) | GCGTGTATGGCAACATGAAC |
| **B17050** | CNA1 5'-flanking region primer L2 (HYG) | GCACTTTTCGGGGAAATGTGGCAAGAGGTATTTGAACGGG |
| **B17051** | CNA1 3'-flanking region primer R1 (HYG) | CGATGAGGTAGTGCAAGTTAGTACACCCAATCTAGGATCT |
| **B17052** | CNA1 3'-flanking region primer R2 (HYG) | CGGACTCGTAGAATCGAAGC |
| **B17053** | CNA1 5'-screening primer SO (HYG) | CTGCCATTGACGAATTGTTG |
| **B17055** | CNA1 3'-sreening primer SO2 (HYG) | CAAGAACACCTGCTCATCCA |
| **B17054** | CNA1 Southern blot probe primer PO (HYG) | TCATTTTGAAGGCCTAGTGTCA |
| **B17100** | *CNA1* sequencing primer 1 | ATGTCTGTGAGAAAAAAGTT |
| **B17101** | *CNA1* sequencing primer 2 | CCACCAGGCTTCGAGAATTA |
| **B17102** | *CNA1* sequencing primer 3 | TCGAATGTACAAGCGCACGA |
| **B17326** | Screening primer for *CNA1* complementation | CTCTCGTGCCGTATTTGACA |
| **B21391** | *CNA1* 5'-flanking region primer L1 (B11220, B11221, B11245) | GGCCACGGTGAAGTATGTTT |
| **B21392** | *CNA1* 5'-flanking region primer L2 (B11220, B11221) | TTCAGATCCACTAGTTCTAGGCAAGAGGTATTTGAACGGG |
| **B21393** | *CNA1* 3'-flanking region primer R1 (B11220) | TTTCCAGAATTTCACTCTTATTACACCCGATCTAGGATCT |
| **B21394** | *CNA1* 3'-flanking region primer R2 (B11220, B11221) | CCCAGAAAGATGCGGTAGAC |
| **B17783** | *CNA1* 3'-flanking region primer L2 (NEO) (B11221) | gtattctgggcctccatgtcGCAAGAGGTATTTGAACGGG |
| **B17784** | *CNA1* 3'-flanking region primer R1 (NEO) (B11221) | gaatgctggtcgctatactgGTACACCCAATCTAGGATCT |

| | | |
|---|---|---|
| **B21398** | *CNA1* 3'-flanking region primer R2 (B11245) | TCGTGATAGCAAATCGCAGA |
| **B21476** | *CNA1* Internal screening primer LP (B11245) | TGCAAGCACTTGACCGATTA |
| **B21477** | *CNA1* Internal screening primer RP(B11245) | GGGAGGACCACGTAAAGACA |
| **B21798** | *CNA1* 5'-flanking region primer L2 (NEO) (B11245) | gtattctgggcctccatgtcGCAAGAGGTATTTGAACAGG |
| **B21799** | *CNA1* 3'-flanking region primer R1 (NEO) (B11245) | gaatgctggtcgctatactgGAAAATCACCGACATGTTGG |
| **B13364** | *CNA1* qRT-PCR primer 1 | GAATGTACAAGCGCACGAAA |
| **B13365** | *CNA1* qRT-PCR primer 2 | GCACTGCAGCCTTGTTGTTA |
| **B13366** | *CNB1* qRT-PCR primer 1 | GAGTGGGGCCATTGATAAGA |
| **B13367** | *CNB1* qRT-PCR primer 2 | ATCGTACCGTCGTGATCCTC |
| **B17348** | Screening primer for *CNB1* complementation | GAGCCGACAGTGAACTCCTC |
| **B17349** | Screening primer for *CRZ1* complementation | CTACCGACTCTGAGGCAAGG |
| **B16985** | *CNB1* sequencing primer 1 | AACCCAAGAAAGCGTTTGGG |
| **B16986** | *CNB1* sequencing primer 2 | ATATGATATCGACAGAGATG |
| **B16991** | *CRZ1* sequencing primer 1 | ATATCTTTCTACGCTTGCAG |
| **B16992** | *CRZ1* sequencing primer 2 | ATCCTAACCCTTCTGGGCTC |
| **B16993** | *CRZ1* sequencing primer 3 | TCCGCAAGTAACAGCTTGAA |
| **B13492** | *CRZ1* qRT-PCR primer 1 | GTGTCGGTGAAAATGCTGTG |
| **B13493** | *CRZ1* qRT-PCR primer 2 | ATGGCAGGCATACAAAGAGG |
| **B17333** | *CRZ2* 5'-flanking region primer L1 | TGGGGGTTTCTTGAAAAGTG |
| **B17334** | *CRZ2* 5'-flanking region primer L2 | TTCAGATCCACTAGTTCTAGGGGAGGAATTTGGGTGGATG |
| **B17335** | *CRZ2* 3'-flanking region primer R1 | TTTCCAGAATTTCACTCTTAACACCGTGAGGTGACACCGG |
| **B17336** | *CRZ2* 3'-flanking region primer R2 | TAGACGGCTGAAGGCAAACT |
| **B17337** | *CRZ2* 5'-screening primer SO | CGTCTTTTGTCAGCCGTGTA |
| **B17339** | *CRZ2* 3'-sreening primer SO2 | AGGTTTGCCTGACCTTGTTG |
| **B17338** | *CRZ2* Southern blot probe primer PO | TGGTGAGCTACCGAAAAACC |
| **B17340** | *CRZ2* Internal screening primer LP | TCAGCAAGTGCAACAAGTCC |
| **B17341** | *CRZ2* Internal screening primer RP | CGTCCAGGTCTCTTCTTTGC |
| **B17755** | *CRZ2* 5'-flanking region primer L1 (NEO) | TGGGGGTTTCTTGAAAAGTG |
| **B17756** | *CRZ2* 5'-flanking region primer L2 (NEO) | gtattctgggcctccatgtcGGGAGGAATTTGGGTGGATG |
| **B17757** | *CRZ2* 3'-flanking region primer R1 (NEO) | gaatgctggtcgctatactgACACCGTGAGGTGACACCGG |
| **B17758** | *CRZ2* 3'-flanking region primer R2 (NEO) | TAGACGGCTGAAGGCAAACT |
| **B17759** | *CRZ2* 5'-screening primer SO (NEO) | CGTCTTTTGTCAGCCGTGTA |
| **B17761** | *CRZ2* 3'-sreening primer SO2 (NEO) | AGGTTTGCCTGACCTTGTTG |
| **B17760** | *CRZ2* Southern blot probe primer PO (NEO) | CCATCCTTTCCCTGATTTGA |
| **B20346** | *CRZ2* qRT-PCR primer 1 | TCAGCAAGTGCAACAAGTCC |
| **B20347** | *CRZ2* qRT-PCR primer 2 | CTGACTGCGTCCTTGGAAAT |

| | | |
|---|---|---|
| **B11749** | *ACT1* qRT-PCR primer 1 | TTGCTCCTGAAGAACACCCT |
| **B11750** | *ACT1* qRT-PCR primer 2 | GCAGGAACGTTGAAGGTCTC |
| **B12554** | *ERG11* qRT-PCR primer 1 | TGCCCATCGTCTACAACCTT |
| **B12555** | *ERG11* qRT-PCR primer 2 | TCTCTCTGCACAGCTCGAAA |
| **B12909** | *ERG6* qRT-PCR primer 1 | AGAGACCAAGAGTTCGCCAA |
| **B12910** | *ERG6* qRT-PCR primer 2 | TTAGCAACGTCAGCAGCATC |
| **B12660** | *FKS1* qRT-PCR primer 1 | CGAAGAACACGGTCAGGACA |
| **B12661** | *FKS1* qRT-PCR primer 2 | CCTCAGGGGTCAAGACGTTC |
| **B17569** | *FKS2* qRT-PCR primer 1 | AACTCCGATGACGTTGAACC |
| **B17570** | *FKS2* qRT-PCR primer 2 | TTGAGCCTCGGAGTTGTCTT |
| **B13289** | *CHS1* qRT-PCR primer 1 | GCCTGAAAGTATCCCGGAGT |
| **B13290** | *CHS1* qRT-PCR primer 2 | CCAAATCCTAGTCGCATGCC |
| **B13291** | *CHS2* qRT-PCR primer 1 | CGGCAGAACAGTTTACGACC |
| **B13292** | *CHS2* qRT-PCR primer 2 | GGGCTTCTGTCTCACCTCTT |
| **B13293** | *CHS3* qRT-PCR primer 1 | GGAGAGAGAAGATGGGGCTC |
| **B13294** | *CHS3* qRT-PCR primer 2 | GTGGTATTGTGGCAAACGGT |
| **B13295** | *CHS4* qRT-PCR primer 1 | GGGTGAAGTTGTCGAATCGG |
| **B13296** | *CHS4* qRT-PCR primer 2 | GGCACAGATGGAGAGCATTG |
| **B13297** | *CHS5* qRT-PCR primer 1 | GTGGGTAAACTCGATGCGTC |
| **B13298** | *CHS5* qRT-PCR primer 2 | TAACAATCGAGCCGGCTTTG |
| **B13299** | *CHS6* qRT-PCR primer 1 | TGCACACATACATTGGCGAG |
| **B13300** | *CHS6* qRT-PCR primer 2 | CTGGATCGTCTGCACACATG |
| **B13301** | *CHS7* qRT-PCR primer 1 | GGTATTGTGGGTGCCTTGTG |
| **B13302** | *CHS7* qRT-PCR primer 2 | ATACCCACATCGACCGAGTC |
| **B13305** | *CDA2* qRT-PCR primer 1 | CAGCTCCAGTGGTCGATTTG |
| **B13306** | *CDA2* qRT-PCR primer 2 | TTGCACGAACTCTGTTGTCG |
| **B20922** | CLP for *CRZ1-mCherry* complementation | GGGCCCATCCATGGTCCATTGAGGGT |
| **B19089** | CRP for *CRZ1-mCherry* complementation | GCGGCCGCGTCTGGTGGCGAGATGGCGA |

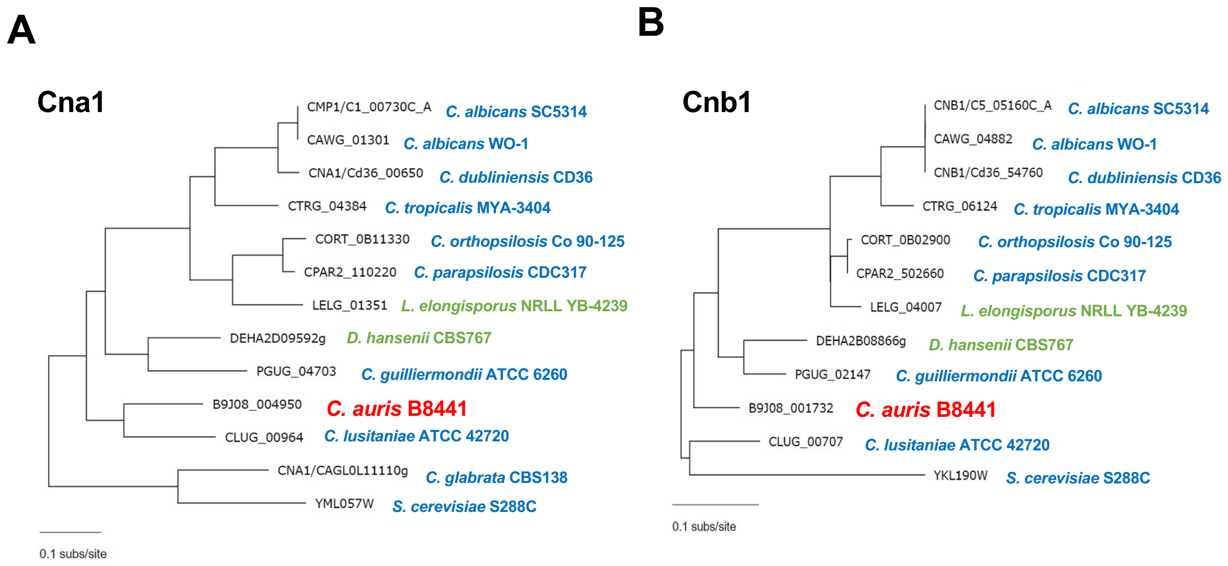


**S1 Fig. Phylogenetic analysis of Cna1 and Cnb1.** Phylogenetic trees for fungal orthologs for Cna1 (A) and Cnb1 (B) across various fungal species were constructed using data from the *Candida* Genome Database (http://www.candidagenome.org/).

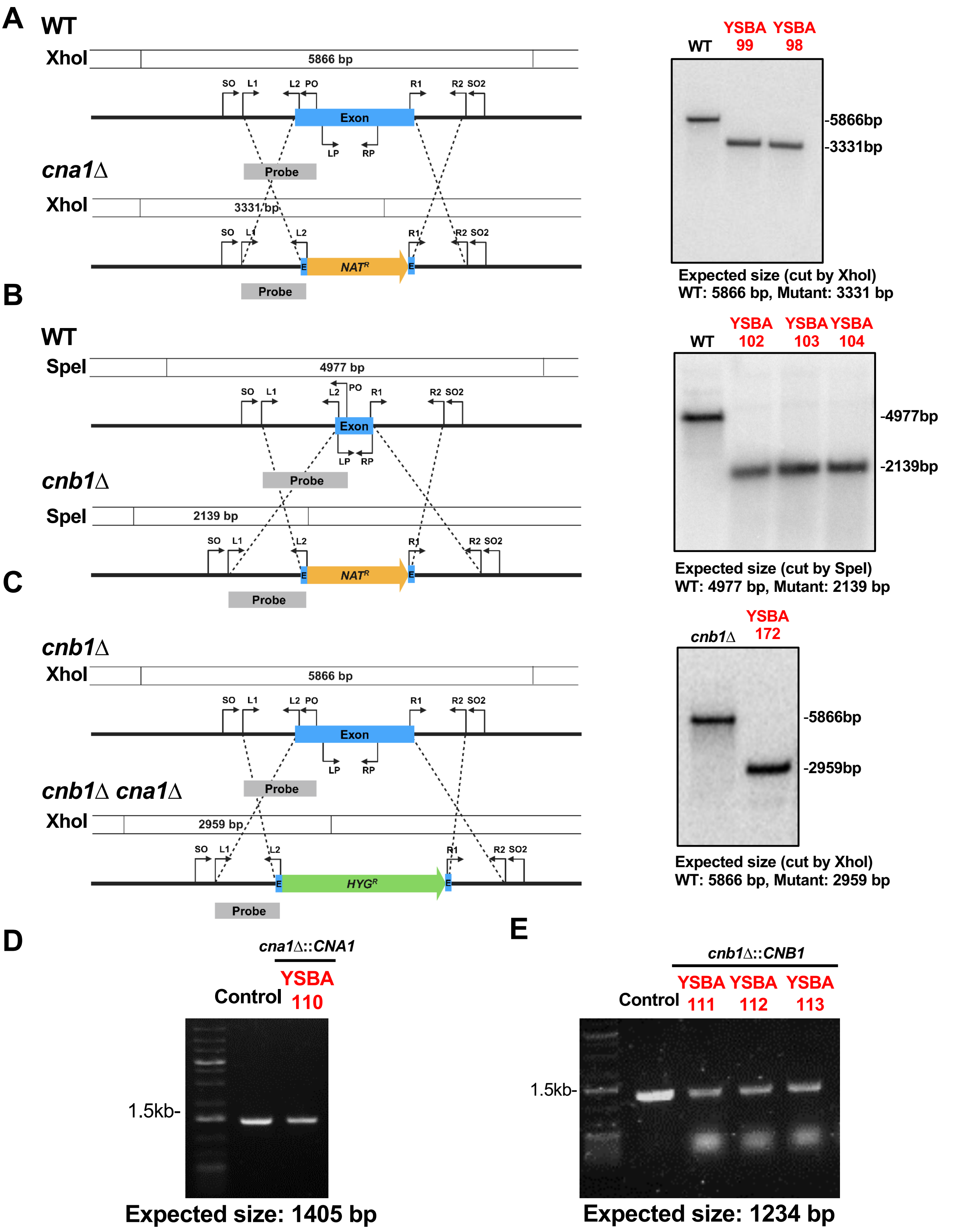


**S2 Fig. Construction and verification of calcineurin gene deletion mutants and complemented strain in *C. auris*.** (A-C) Schematic representation of the homologous recombination strategies used to delete *CNA1*, *CNB1*, and both genes (left panels). Southern blot analyses confirm the successful deletion of the target genes (right panels). (D, E) Verification of the constructed complemented strains by diagnostic PCR.

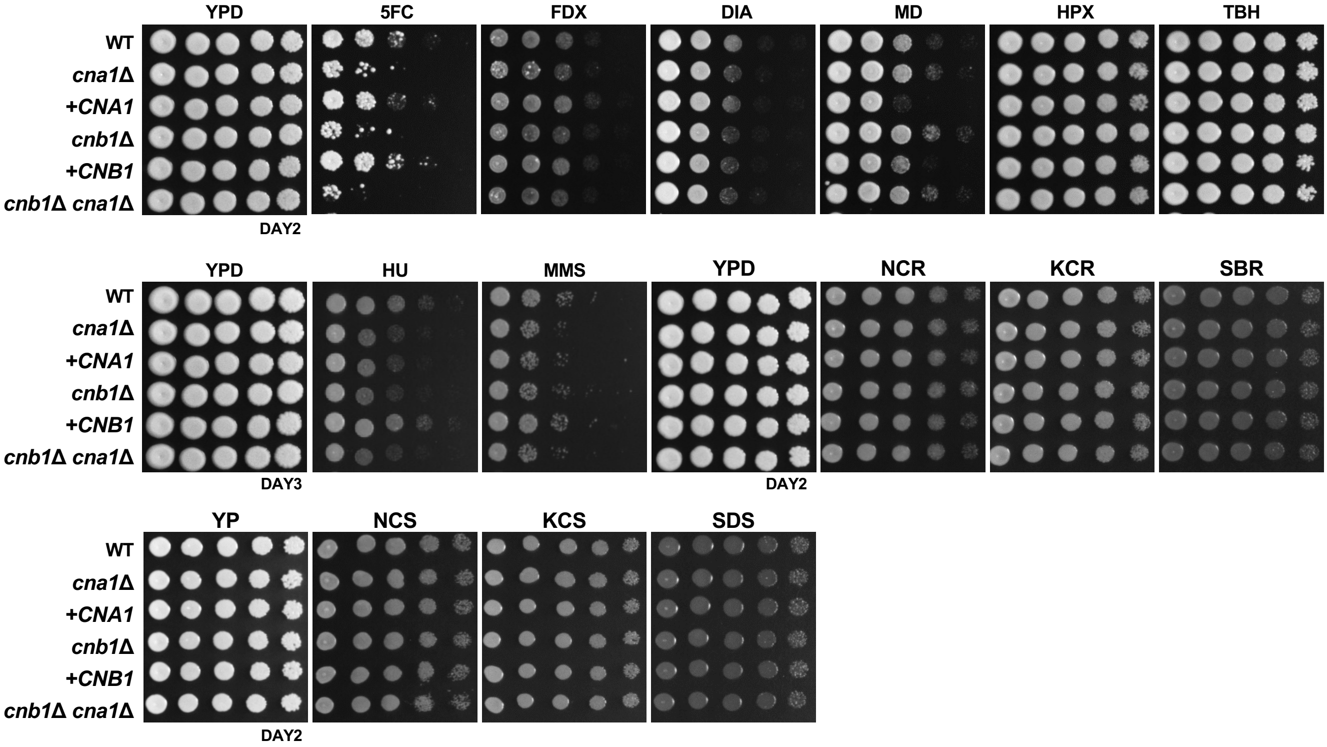


**S3 Fig. Qualitative spot assays showing stress response of calcineurin mutants.** WT (B8441), *cna1*Δ (YSBA99), *cna1*Δ::*CNA1* (*+CNA1*; YSBA110), *cnb1*Δ (YSBA102), *cnb1*Δ::*CNB1* (*+CNB1*; YSBA113), and *cnb1*Δ *cna1*Δ (YSBA172) strains were spotted on YPD medium supplemented with stressors such as 50 µg/ml 5FC, 3 µg/ml FDX, 2.5 mM DIA, 0.06 mM MD, 10m M HP, 2.1 mM *tert*-butyl hydroperoxide (TBH), 150 mM hydroxyurea (HU), 0.03% MMS, 1.5 M NaCl, 1.5 M KCl, or 2 M sorbitol. WT and mutant strains were spotted on YP medium supplemented with 1 M NaCl, 1 M KCl, or 2 M Sorbitol. Plates were incubated for 2 or 3 days. Abbreviations: 5FC, 5-flucytosine; FDX, fludioxonil; DIA, diamide; MD, menadione; HPX, hydrogen peroxide; TBH, *tert*-butyl hydroperoxide; HU, hydroxyurea; MMS, methyl methanesulphonate; KCR, YPD + KCl; NCR, YPD + NaCl; SBR, YPD + sorbitol; KCS, YP + KCl; NCS, YP + NaCl; SBS, YP + sorbitol.

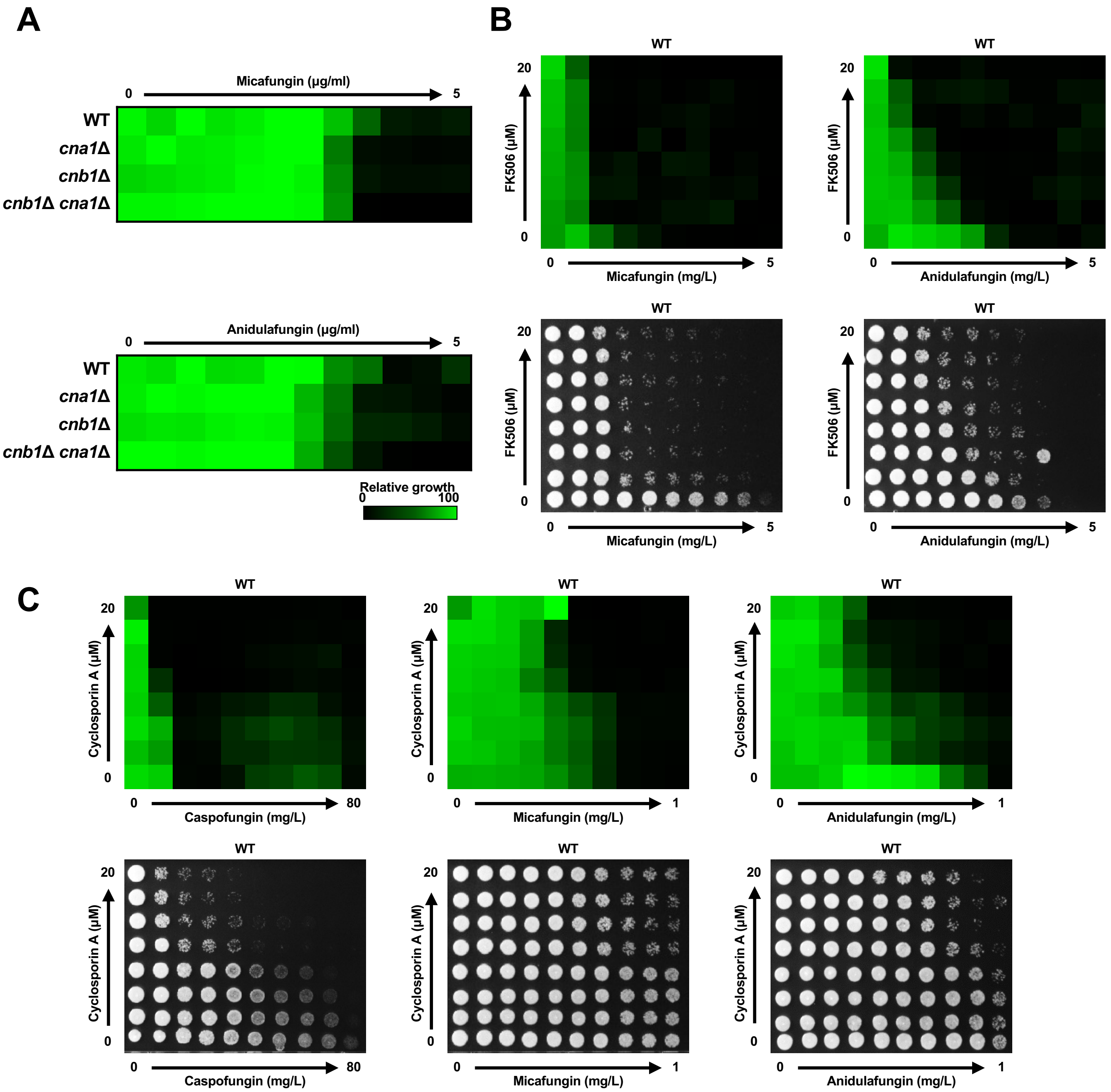


**S4 Fig. MIC tests of echinocandins and checkerboard assays with FK506, cyclosporin A, and echinocandins.** (A) EUCAST MIC test results for micafungin (MIF) and anidulafungin (ANF) in the WT (B8441), *cna1*Δ (YSBA99), and *cnb1*Δ (YSBA102) strains. (B) Checkerboard assay results showing the interaction between MIF or ANF and FK506 in the WT strain (B8441). (C) Checkerboard assay results of echinocandins with cyclosporin A in the WT strain (B8441).

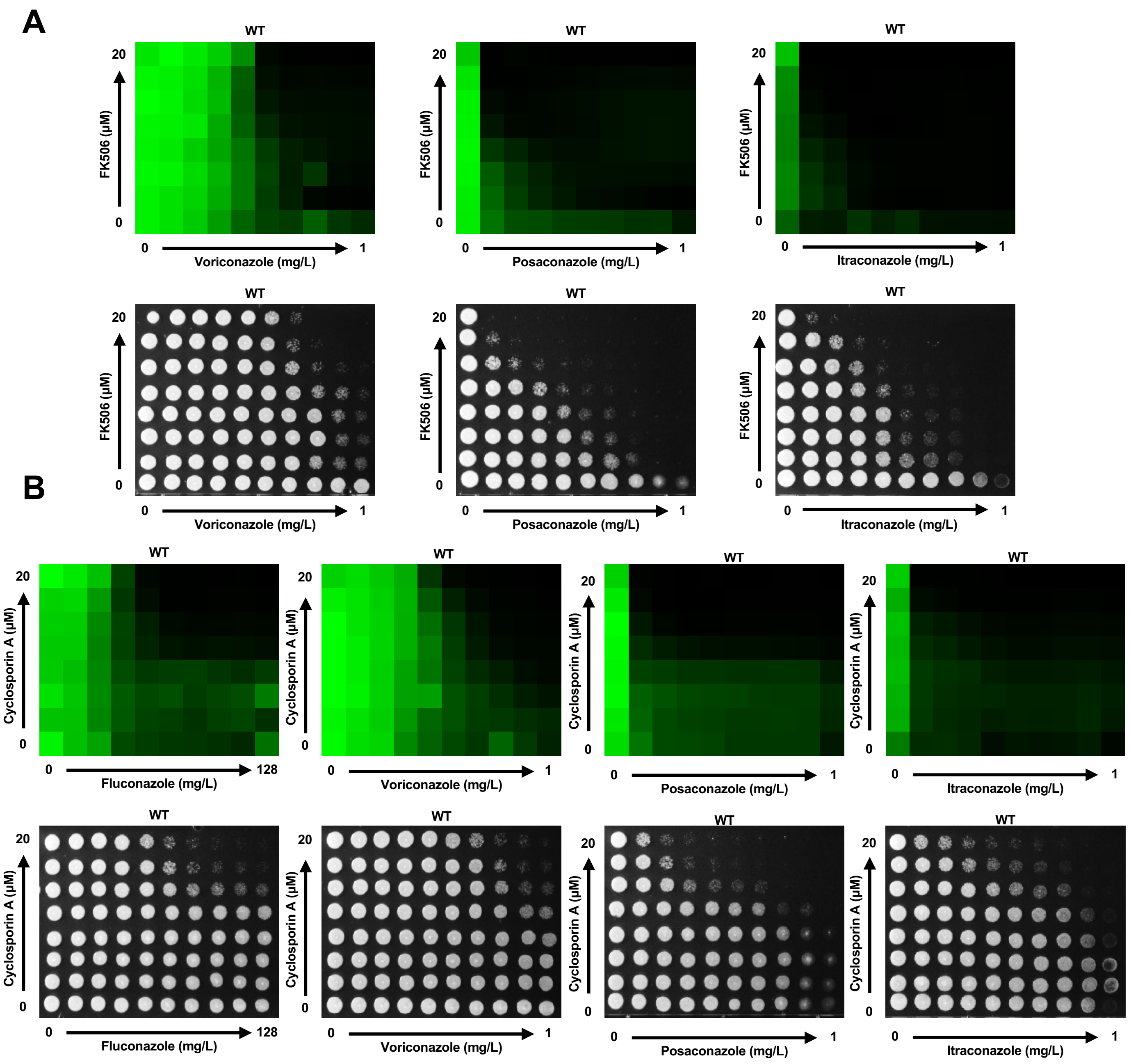


**S5 Fig. Checkerboard assays with FK506, cyclosporin A, and azoles.** (A) Checkerboard assay results showing the interaction between voriconazole (VRC), posaconazole (PSC), or itraconazole (ITC) and FK506 in the WT strain (B8441). (B) Checkerboard assay results depicting the interaction of fluconazole (FLC), VRC, PSC, or ITC with cyclosporin A in the WT strain (B8441).

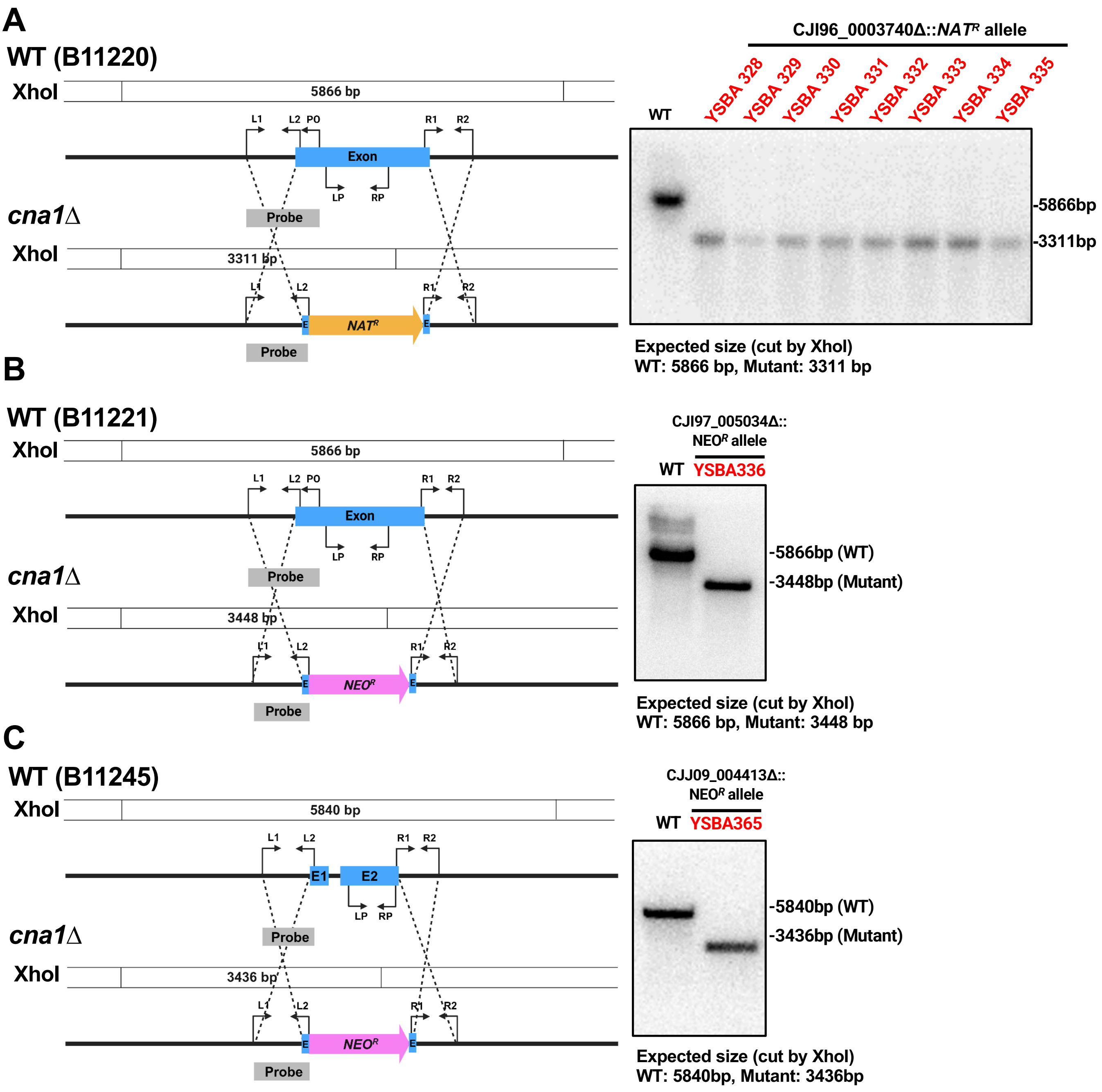


**S6 Fig. Construction and verification of *cna1*Δ mutants in different clades of *C. auris*.** (A-C) Diagrams illustrating the homologous recombination strategies used to delete *CNA1* in clade II B11220 (A), clade III B11221 (B), and clade IV B11245 (C) strains (left panels). Successful deletion of *CNA1* was confirmed via Southern blot analysis (right panels).

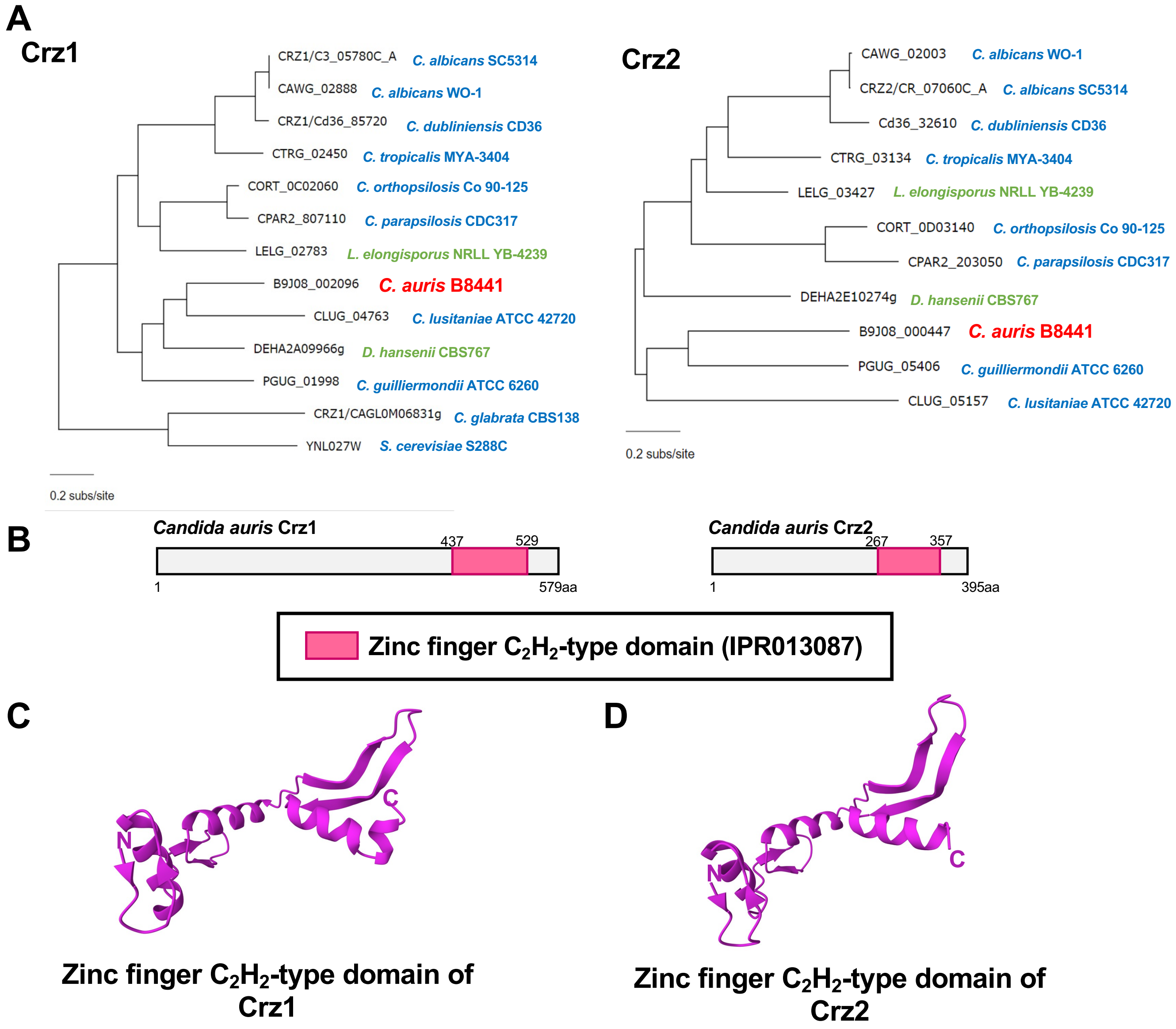


**S7 Fig. Identification of potential downstream transcription factors of calcineurin.** (A) Phylogenetic analysis of Crz1 and Crz2 orthologs across various fungal species. (B) Protein domain analysis of Crz1 and Crz2 in *C. auris*. (C and D) Predicted structure of zinc finger C2H2-type domain of Crz1 and Crz2, generated using AlphaFold2 (ColabFold v1.5.2).

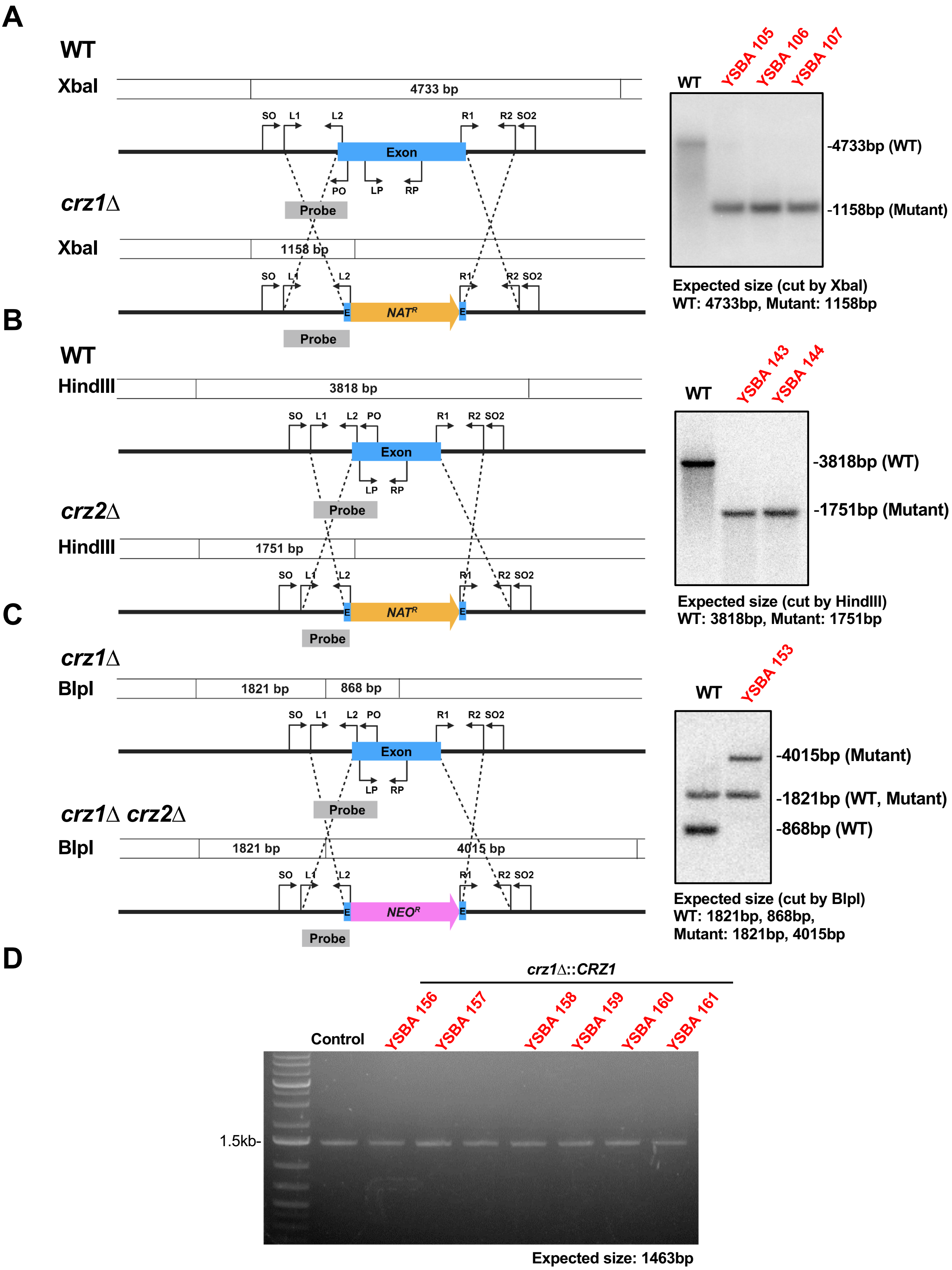


**S8 Fig. Construction and verification of *crz1Δ*, *crz2Δ*, and *crz1Δ crz2Δ* mutants and complemented strains in *C. auris*.** (A-C) Diagrams illustrating the homologous recombination strategies used to delete *CRZ1* (A), *CRZ2* (B), and both genes (C) in the B8441 strain (left panels). Successful gene deletion was confirmed via Southern blot analyses (right panels). (D) Verification of the *crz1Δ::CRZZ1* complemented strains by diagnostic PCR.

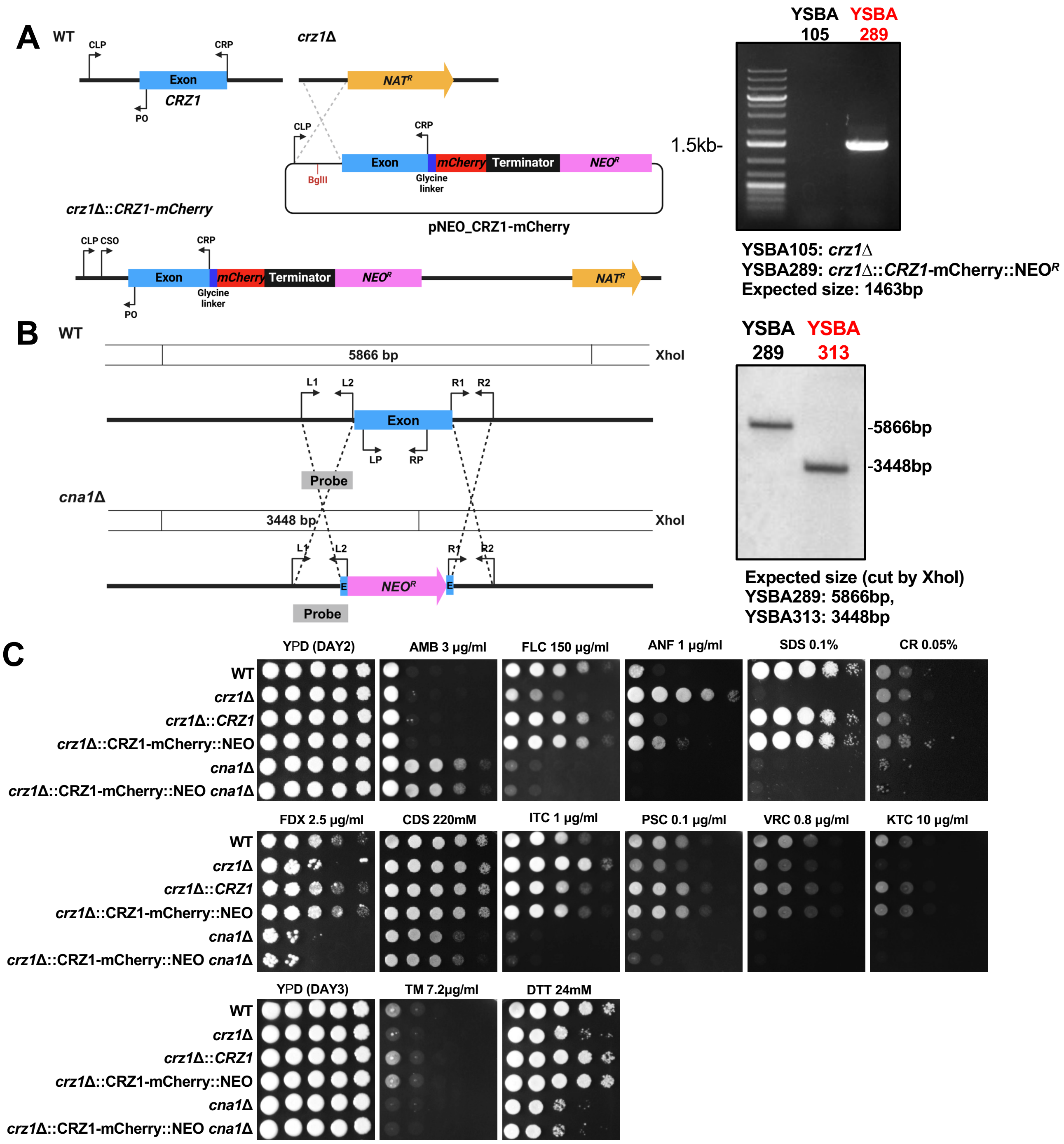


**S9 Fig. Construction and verification of mCherry-tagged Crz1 strains in *C. auris*.** (A) Diagrams illustrating the homologous recombination strategies used to integrate the *CRZ1-mCherry* allele in the *crz1*Δ mutant (left panel). Diagnostic PCR confirmed the targeted integration of *CRZ1-mCherry* in the tagged strains. (B) Diagrams illustrating the homologous recombination strategies used to delete *CNA1* in the *CRZ1-mCherry* stain (left panel). Disruption of the *CNA1* in the *CRZ1-mCherry* strain was validated through Southern blot analysis. (C) Qualitative spot assays showing the phenotypes of WT (B8441), *crz1*Δ (YSBA105), *crz1*Δ::*CRZ1* (YSBA158), *crz1*Δ::*CRZ1-mCherry* (YSBA289), *cna1*Δ (YSBA99), and *crz1*Δ::*CRZ1-mCherry cna1*Δ (YSBA313) strains. WT and mutant strains were spotted on YPD medium supplemented with stressors, including 3 μg/ml amphotericin B (AMB), 150 μg/ml fluconazole (FLC), 1 μg/ml anidulafungin (ANF), 0.1% SDS, 0.05% Congo red (CR), 2.5 μg/ml fludioxonil (FDX), 220 mM $CdSO_4$ (CDS), 1 μg/ml itraconazole (ITC), 0.1 μg/ml posaconazole (PSC), 0.8 μg/ml voriconazole (VRC), 10 μg/ml ketoconazole (KTC), 7.2 μg/ml tunicamycin (TM), or 24 mM DTT. Plates were incubated for 2 or 3 days.

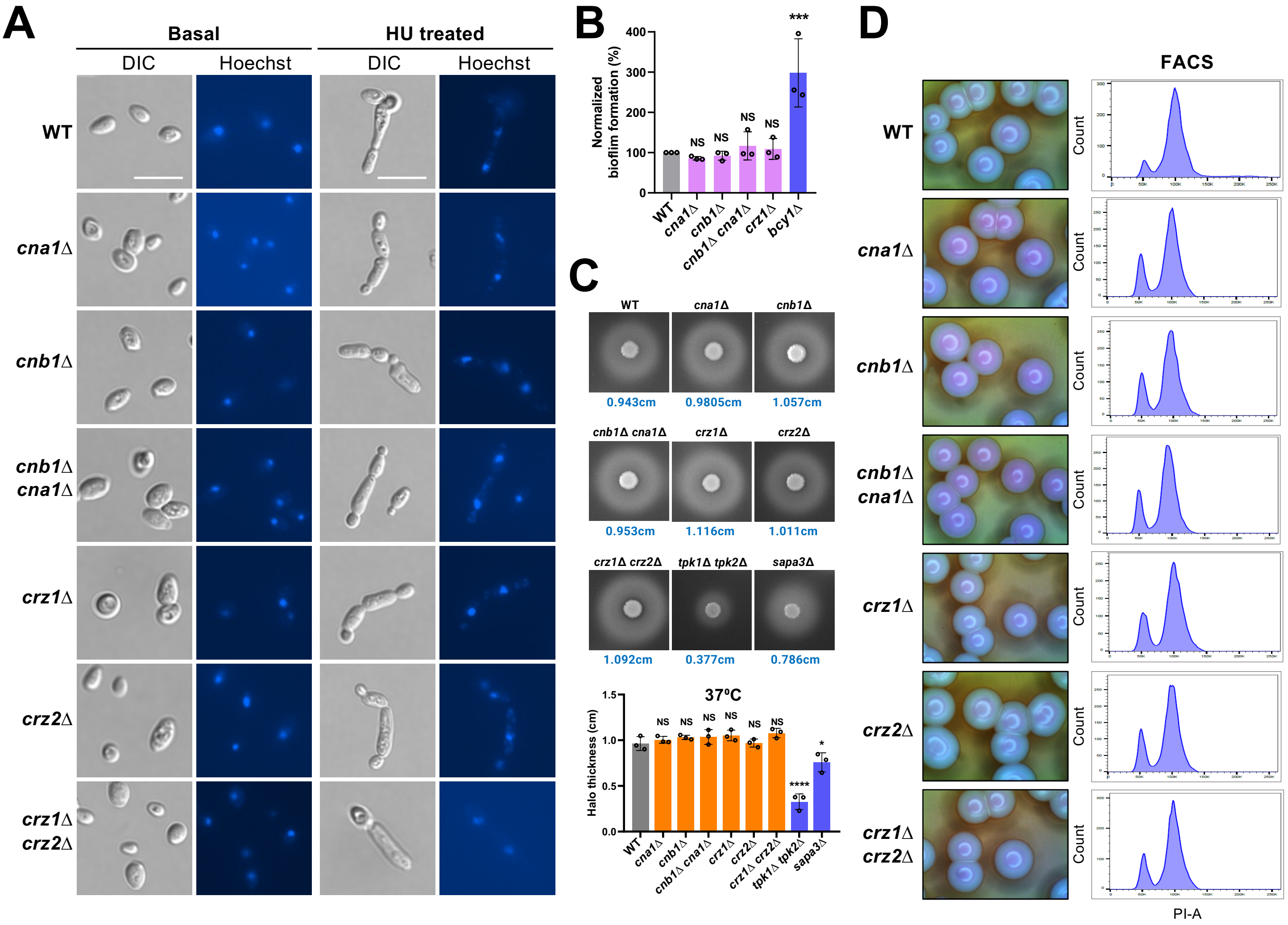


**S10 Fig. Calcineurin is dispensable for morphogenesis, biofilm formation, SAP activity, and ploidy switching in *C. auris*.** (A) Hydroxyurea (HU)-mediated filamentous growth. Indicated WT and mutant cells were cultured overnight in YPD medium at 30°C and then subcultured into fresh YPD medium to an OD$_{600}$ of 0.8. The cultures were treated with 100 mM HU in YPD broth and incubated for 24 h at 30°C. The cells were fixed using 10% formalin and stained with Hoechst solution. Representative microscopy images of each strain are shown. (B) Biofilm formation assay. Biofilm formation by WT and mutant strains was assessed using crystal violet staining. The absorbance of the destaining solution for each strain was measured at 595 nm. The *bcy1*Δ mutant was used as a positive control. (C) Secreted aspartyl protease (SAP) activity assay. WT and mutant strains were cultured overnight, washed twice with dH$_2$O, resuspended in dH$_2$O, spotted (3 µL) onto solid YCB-BSA medium. The plates were incubated for 3 days at 37°C, and the halo diameter was measured to determine SAP activity. Experiments was biologically replicated three times. Statistical significance was assessed using one-way ANOVA with Bonferroni's multiple-comparison test (*, $P < 0.05$; **, $P < 0.01$; ***, $P < 0.001$; ****, $P < 0.0001$; NS, not significant). The *tpk1*Δ *tpk2*Δ and *sapa3*Δ mutants were used as negative controls. (D) Ploidy switching assessment. Approximately 100 cells from WT and mutant strains were plated on YPD medium containing 5 µg/ml phloxine B, incubated at 25°C for 14 days, and photographed. Cells isolated from phloxine B-containing plates were cultured in liquid YPD medium at 30°C for 48 h, fixed, photographed, and analyzed by FACS.
